# A mutant ASXL1-EHMT complex contributes to heterochromatin dysfunction in clonal hematopoiesis and chronic monomyelocytic leukemia

**DOI:** 10.1101/2024.01.30.578015

**Authors:** Zhen Dong, Hugo Sepulveda, Leo J. Arteaga-Vazquez, Chad Blouin, Jenna Fernandez, Moritz Binder, Wen-Chien Chou, Hwei-Fang Tien, Mrinal Patnaik, Geoffrey J Faulkner, Samuel A Myers, Anjana Rao

**Affiliations:** Department of Signaling and Gene Expression, La Jolla Institute for Allergy and Immunology, La Jolla, California 92037; Sanford Consortium for Regenerative Medicine, La Jolla, California 92037; Department of Pharmacology, University of California, San Diego, California 92161; Moores Cancer Center, University of California, San Diego, California 92037; Division of Hematology, Department of Internal Medicine, Mayo Clinic, Rochester, Minnesota 55905; Division of Hematology, Department of Internal Medicine, National Taiwan University Hospital, No. 7, Chung-Shan S Rd, Taipei 10002, Taiwan; Mater Research Institute - University of Queensland, TRI Building, Woolloongabba, QLD, 4102, Australia; Queensland Brain Institute, University of Queensland, Brisbane QLD 4072, Australia

**Keywords:** Clonal Hematopoiesis, CMML, ASXL1/BAP1, heterochromatin, EHMT1/2 (GLP/G9a)

## Abstract

*ASXL1* is one of the three most frequently mutated genes in age-related clonal hematopoiesis (CH), alongside *DNMT3A* and *TET2*. CH can progress to myeloid malignancies including chronic monomyelocytic leukemia (CMML), and is also strongly associated with inflammatory cardiovascular disease and all-cause mortality in humans. DNMT3A and TET2 regulate DNA methylation and demethylation pathways respectively, and loss-of-function mutations in these genes reduce DNA methylation in heterochromatin, allowing de-repression of silenced elements in heterochromatin. In contrast, the mechanisms that connect mutant ASXL1 and CH are not yet fully understood. CH/CMML-associated *ASXL1* mutations encode C-terminally truncated proteins that enhance the deubiquitinase activity of the ASXL-BAP1 “PR-DUB” deubiquitinase complex, which removes mono-ubiquitin from H2AK119Ub. Here we show that ASXL1 mutant proteins interact with the EHMT1-EHMT2 methyltransferase complex, which generates H3K9me1 and me2, the latter a repressive modification in constitutive heterochromatin. Compared to cells from age-matched wildtype mice, we found that expanded myeloid cells from old (≥18-month-old) *Asxl1tm/+* mice, a heterozygous knock-in mouse model of CH, display genome-wide decreases of H3K9me2, H3K9me3 and H2AK119Ub as well as an associated increase in expression of transposable elements (TEs) and satellite repeats. Increased TE expression was also observed in monocytes from *ASXL1*-mutant CMML patients compared to monocytes from healthy controls. Our data suggest that mutant ASXL1 proteins compromise the integrity of both constitutive and facultative heterochromatin in an age-dependent manner, by reducing the levels of H3K9me2/3 and H2AK119Ub. This increase in TE expression correlated with increased expression of nearby genes, including many interferon-inducible (inflammation-associated) genes (ISGs).

**Significance Statement:** Age-related clonal hematopoiesis (CH) is a premalignant condition associated with inflammatory cardiovascular disease. *ASXL1* mutations are very frequent in CH. We show that ASXL1 interacts with EHMT1 and EHMT2, H3K9 methyltransferases that deposit H3K9me1 and me2. Using a mouse model of mutant *ASXL1* to recapitulate CH, we found that old ASXL1-mutant mice showed marked expansion of myeloid cells in bone marrow, with decreased H3K9me2/3 and increased expression of transposable elements (TEs) in heterochromatin. In humans, ASXL1-mutant CH progresses to chronic monomyelocytic leukemia (CMML); CMML patient samples showed striking upregulation of many TE families, suggesting that ASXL1 mutations compromise heterochromatin integrity, hence causing derepression of TEs. Targeting heterochromatin-associated proteins and TEs might counter the progression of CH, CMML and other myeloid malignancies.

## Introduction

ASXL1, ASXL2 and ASXL3, the human homologues of the *Drosophila Asx* (*additional sex combs*) gene, are obligate regulatory components of the PR-DUB BAP1 deubiquitinase complex, which removes ubiquitin from H2AK119Ub (1–3). Somatic mutations in *ASXL1* are frequent in clonal hematopoiesis (CH), an age-associated condition that can progress to myeloid malignancies including myelodysplastic syndrome (MDS), chronic monomyelocytic leukemia (CMML) and acute myeloid leukemia (AML) (4, 5), and germline mutations of *ASXL1* and *ASXL3* have been implicated in the neurodevelopmental disorders Bohring-Opitz and Bainbridge-Ropers syndromes respectively (6).

We previously showed that CH/CMML-associated ASXL1-mutant proteins enhance BAP1 activity (7). Here we show that C-terminally truncated ASXL1 mutant proteins also interact with the EHMT1-EHMT2 (GLP-G9a) H3K9 methyltransferase complex, which is responsible for generation of the H3K9me1 and me2 marks (8, 9). Using a knock-in mouse model of CH (*Asxl1tm/+* mice)(10), we show that expanded myeloid cells from old mice (≥18-month-old) display genome-wide decreases of H3K9me2, H3K9me3 and H2AK119Ub and substantial changes in gene expression compared to cells from age-matched wildtype mice; cells from young (4-month-old) mice showed no appreciable differences. Compared to control LK (lineage-negative, c-Kit+) cell populations enriched for hematopoietic stem/precursor cells (HSPC), LK cells from aged *Asxl1tm/+* mice showed increased expression of transposable elements (TEs) and satellite repeats, consistent with perturbed heterochromatin integrity. Increased TE and gene expression in heterochromatin were also observed in bone marrow samples from *ASXL1*-mutant CMML patients compared to healthy controls. In a small number of cases in which individual TEs could be mapped uniquely to the genome, increased TE expression was shown to alter the expression of nearby genes in *cis*, and was associated with altered expression of inflammation-associated, interferon-inducible genes (ISGs).

In summary, we find that mutant ASXL1 compromises heterochromatin integrity in an age-dependent manner by decreasing H3K9me2/3 levels in constitutive heterochromatin and H2AK119Ub levels in facultative heterochromatin. We propose that the top three mutations in clonal hematopoiesis share the common feature of promoting heterochromatin dysfunction, through reduction of heterochromatic DNA methylation in *TET2-* and *DNMT3A-* mutant cells (11) or decreased levels of H3K9me2/3 and H2AK119Ub histone modification in *ASXL1*-mutant cells.

## Results

### Cancer-associated ASXL1 mutant proteins bind heterochromatin-associated proteins

The heterozygous *ASXL1* and *ASXL3* mutations observed in CH, CMML, Bohring-Opitz and Bainbridge-Ropers syndrome invariably encode C-terminally truncated proteins (6, 12, 13). Essentially all frameshift and nonsense mutations occur in the large penultimate exons of *ASXL* genes, thus all ASXL-mutant proteins retain the DEUBAD deubiquitinase adapter domain (aa 240-390 of human ASXL1) (shown for *ASXL1* in **Fig. 1A**, *top* and ***SI Appendix,* Fig. S1**). The frameshift mutation c.1934dupG; p.G646WfsX12 (here termed ASXL1G646fs) is one of the most common, both in pre-neoplastic CH samples and in myeloid neoplasms (MPN) including the myelodysplastic/ myeloproliferative overlap neoplasm CMML (12, 13) (***SI Appendix*, Fig. S1; Fig. 1A**, *top*).

**Figure 1.**
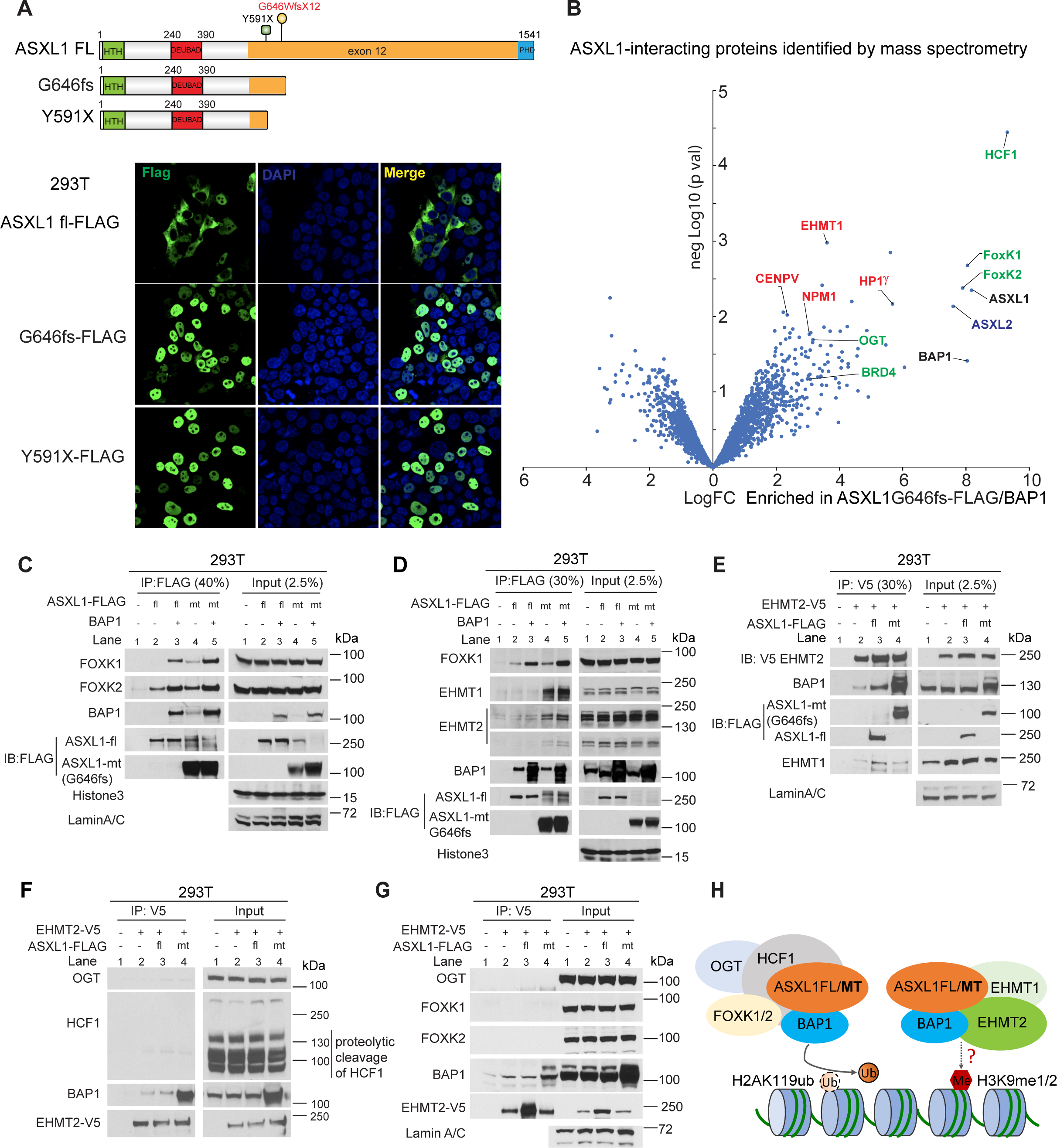
C-terminally truncated ASXL1 proteins reside in the nucleus and recruit novel epigenetic regulators. **A.** *Top*, Diagram of full-length (fl) ASXL1 and two C-terminally truncated ASXL1 mutant proteins encoded by ASXL1 mutations associated with CH/CMML (also see *SI Appendix*, **Fig. S1**). Full-length ASXL1 contains a predicted DNA binding (HTH) domain at its N-terminus, followed by the DEUBAD deubiquitinase adapter domain and a C-terminal atypical PHD Zinc-finger. A 3xFLAG epitope tag was engineered at the C terminus. *Bottom*, Subcellular localization of C-terminally 3xFLAG-tagged full length ASXL1 and mutant ASXL1G646fs and Y591X proteins in HEK293T cells. **B.** Proteins interacting with mutant ASXL1G646fs. HEK293T cells were co-transfected with ASXL1G646fs-3xFLAG plus BAP1 or with GFP as control, and proteins interacting with mutant ASXL1 were co-immunoprecipitated (co-IP’d) with anti-FLAG antibody and identified by mass spectrometry. The proteins labeled in green in the volcano plot are proteins already reported to interact with ASXL1, while the proteins labeled in red are novel heterochromatin-related regulators identified in this study. **C.** FOXK1 and FOXK2 bind both full-length (fl) ASXL1 and mutant (mt) ASXL1G646fs and the binding is potentiated by BAP1. HEK293T cells were transfected with full-length ASXL1-3xFLAG or mutant ASXL1G646fs-3xFLAG with or without BAP1. Nuclear fractions were isolated and immunoprecipitated with anti-FLAG. ASXL1-bound FOXK1, FOXK2 and BAP1 were detected with antibodies to the endogenous proteins. The bands migrating at the same size as full length ASXL1 in anti-FLAG IPs and input lanes of ASXL1G646fs FLAG (lanes 4, 5) are non-specific bands that only appear when we use the anti-FLAG antibody to IP mutant ASXL1-3xFLAG from cell lysates. They do not appear in IPs of full-length ASXL1-3xFLAG, either using anti-FLAG or an ASXL1 antibody that recognizes an epitope after the DEUBAD domain (not present in truncated mutant ASXL1 Y591X or G646fs). « - » indicates cells transduced with empty vector. **D.** The H3K9 methyltransferases EHMT1 and EHMT2 co-IP more strongly with mutant (mt) ASXL1G646fs compared to full-length (fl) ASXL1. HEK293T cells were transfected with full-length (fl) ASXL1-3xFLAG or mutant (mt) ASXL1G646fs-3xFLAG with or without BAP1. Nuclear fractions were isolated and immunoprecipitated with anti-FLAG. ASXL1-bound FOXK1, EHMT1 and EHMT2 were detected by immunoblotting (IB) using antibodies to the endogenous proteins. The bands at the same size as full length ASXL1 in anti-Flag IPs of mutant ASXL1 G646fs (lanes 4, 5) are the non-specific bands mentioned in **C**. **E.** Reciprocal co-IP with V5-tagged EHMT2. HEK293T cells were co-transfected with V5-EHMT2 and full-length (fl) ASXL1-3xFLAG or mutant (mt) ASXL1G646fs-3xFLAG. Proteins co-IP’ing with EHMT2 were detected with anti-FLAG, anti-EHMT1 and anti-BAP1. **F**, **G**. V5-tagged EHMT2 expressed in HEK293T cells co-IPs with BAP1 but not with OGT and HCF1 (**F**), or with FOXK1 and FOXK2 (**G**). **H**. Diagram of two postulated complexes containing the ASXL1-BAP1 heterodimer, one containing the OGT-HCF1 dimer and possibly also FOXK1 and FOXK2 and the other containing the histone methyltransferases EHMT1 and EHMT2 as homodimers or the heterodimer. BAP1 is thought to tether ASXL1 to nucleosomes through its interaction with the acidic patch, and so may facilitate both the canonical function of the ASXL1-BAP1 PR-DUB complex (the removal of mono-ubiquitin from H2AK119Ub), and the observed decrease in H3K9me2/me3 through mechanisms that are still undefined. FL means the full-length ASXL1 protein, while MT means the mutant ASXL1 protein.

To understand the association of mutant ASXL1 with CH, we identified proteins interacting with mutant ASXL1 by mass spectrometry. Full-length FLAG-tagged ASXL1 ectopically expressed in HEK 293T cells was both cytoplasmic and nuclear (**Fig. 1A**, *top panel*) as previously reported (14), whereas two distinct C-terminal truncation mutants of human ASXL1 (ASXL1G646fs and ASXL1Y591X) were predominantly present in the nucleus (**Fig. 1A**, *middle and bottom panels*). Because our *ASXL1*-mutant mouse model (*Asxl1tm/+*) bears the ASXL1G646fs mutation (10), we used the ASXL1G646fs mutant protein to identify interacting proteins by mass spectrometry.

Mutant ASXL1G646fs is more stable than full-length ASXL1 and the expression of mutant and full-length ASXL1 are both further stabilized by co-expression of BAP1 (14). Previous mass spectrometry experiments were performed after immunoprecipitation of ASXL1 that was ectopically expressed in the absence of BAP1, leading to results that were variable and in some cases contradictory between different experiments (14–20). We reasoned that this might be due to low or variable nuclear expression of ASXL1 proteins that were not stably folded in the absence of BAP1. To avoid this problem and ensure high levels of a properly configured ASXL1G646fs-BAP1 complex in the nucleus, we expressed both BAP1 and FLAG-tagged ASXL1G646fs in HEK293T cells, then co-immunoprecipitated (co-IP’d) interacting proteins from nuclear extracts (***SI Appendix*. Fig. S2A-C**).

Mass spectrometric analysis (**Fig. 1B**) revealed several known ASXL1-BAP1 interactors (HCF1, OGT, FOXK1, FOXK2, BRD4; **Fig. 1B**, *green labels*) (14–20), as well as several proteins whose functions are related to heterochromatin. Among these were the H3K9 methyltransferase EHMT1 (GLP), which forms a tight dimer with EHMT2 (G9a) to generate H3K9me1 and me2 (8, 9); HP1γ (heterochromatin-associated protein-1 gamma), one of three conserved HP1 proteins that bind H3K9me3 (21, 22); and NPM1 (nucleophosmin), a nucleolar protein that is recurrently mutated in AML such that it loses its nucleolar localization signal and becomes localized to the cytoplasm (23). We focused on EHMT1 and EHMT2 which possess enzymatic activity and contribute to heterochromatin integrity by generating H3K9me1 and me2 (21, 22).

Co-IP data showed that ASXL1-FOXK1 and ASXL1-FOXK2 interactions were strengthened if BAP1 was co-expressed (**Fig. 1C and D**, *top panel*, *compare lanes 2, 4 to lanes 3, 5*). In contrast, the interaction of ASXL1G646fs with EHMT1 and EHMT2 was relatively insensitive to co-expression of BAP1 (**Fig. 1D**, *top panels, compare lanes 4, 5*); this was also true for its interactions with NPM1 and HP1γ (***SI Appendix,* Fig S2D, S2E**). To determine if BAP1 was part of the ASXL1-EHMT1-EHMT2 complex, we expressed V5-tagged EHMT2 together with FLAG-tagged mutant ASXL1 or full-length ASXL1 in HEK293T cells. V5-EHMT2 effectively pulled down endogenous EHMT1, endogenous BAP1 and both mutant and full-length ASXL1 from HEK293T cell nuclear lysates (**Fig. 1E**). However, V5-EHMT2 did not pull down the known ASXL1 interactors including HCF1, OGT, FOXK1 and FOXK2 (**Figs. 1F and 1G**), suggesting that the ASXL1/BAP1 heterodimer can form two distinct complexes, with EHMT1 and EHMT2 or with OGT and HCF1 respectively (**Fig. 1H**). It is not yet clear whether FOXK1 and/or FOXK2 are part of the ASXL1-BAP1-OGT-HCF1 complex (**Fig. 1H**), or whether they form a separate, third complex with ASXL1 and BAP1.

K562 cells express both full-length ASXL1 and the ASXL1Y591X mutant protein. Because of the poor sensitivity of the commercially available antibodies to endogenous EHMT1, we were unable to co-IP the endogenous ASXL1Y591X with endogenous EHMT1/2 in K562 cell nuclear lysates. However, antibodies to endogenous EHMT2 did pull down both FLAG-tagged full-length ASXL1 and mutant ASXL1Y591X ectopically and stably expressed in K562 cells (***SI Appendix,* Fig S2F**).

### Age-dependent changes in hematopoietic differentiation and gene expression in ASXL1 mutant mice

We next showed that the heterozygous *Asxl1tm/+* mouse strain is a reliable model of age-related clonal hematopoiesis. The *Asxl1tm* mutation is the mouse correlate of the human G646WfsX12 frame-shift mutation “knocked-in” to the *Asxl1* locus (**Fig. 2A**). The G646WfsX12 mutant protein strongly potentiates the ability of BAP1 to deubiquitinate H2AK119Ub (7). To mirror CH in humans in which *DNMT3A*, *TET2* and *ASXL1* mutations are initially always heterozygous, we examined heterozygous *Asxl1tm/+* knock-in mice. Consistent with the previous report (10), we did not observe any change in the frequencies of hematopoietic cell subsets in bone marrow of young (4-month-old) heterozygous *Asxl1tm1/+* knock-in mice compared to young WT mice(**Figs. 2B-E**, *top panels*); however, we did observe significantly increased frequencies of CD11b^+^ Gr-1^+^ myeloid cells (**Figs. 2B, 2C**, *bottom panels*) and long-term hematopoietic stem cells (LT-HSC, **Figs. 2D, 2E**, *bottom panels*), in bone marrow cells from older *Asxl1tm1/+* mice (18- to 20-month-old) compared to age-matched WT controls. There was also a significant decrease in the frequencies of multipotent myeloid progenitor cells (MPP, **Figs. 2D, 2E**, *bottom panels*) and B220^+^ B cells, pro-B cells (B220^+^CD43^+^), pre-B cells (B220^+^CD43^-^) and Ter119^+^ erythroid cells (stages II and III) (***SI Appendix,* Fig S3A-C**), in old *Asxl1tm/+* mice. The frequencies of common myeloid progenitors (CMP), megakaryocyte-erythrocyte progenitors (MEP) and granulocyte-monocyte progenitors (GMP) were not appreciably altered in either young or old mice (***SI Appendix,* Fig S3D**).

**Figure 2.**
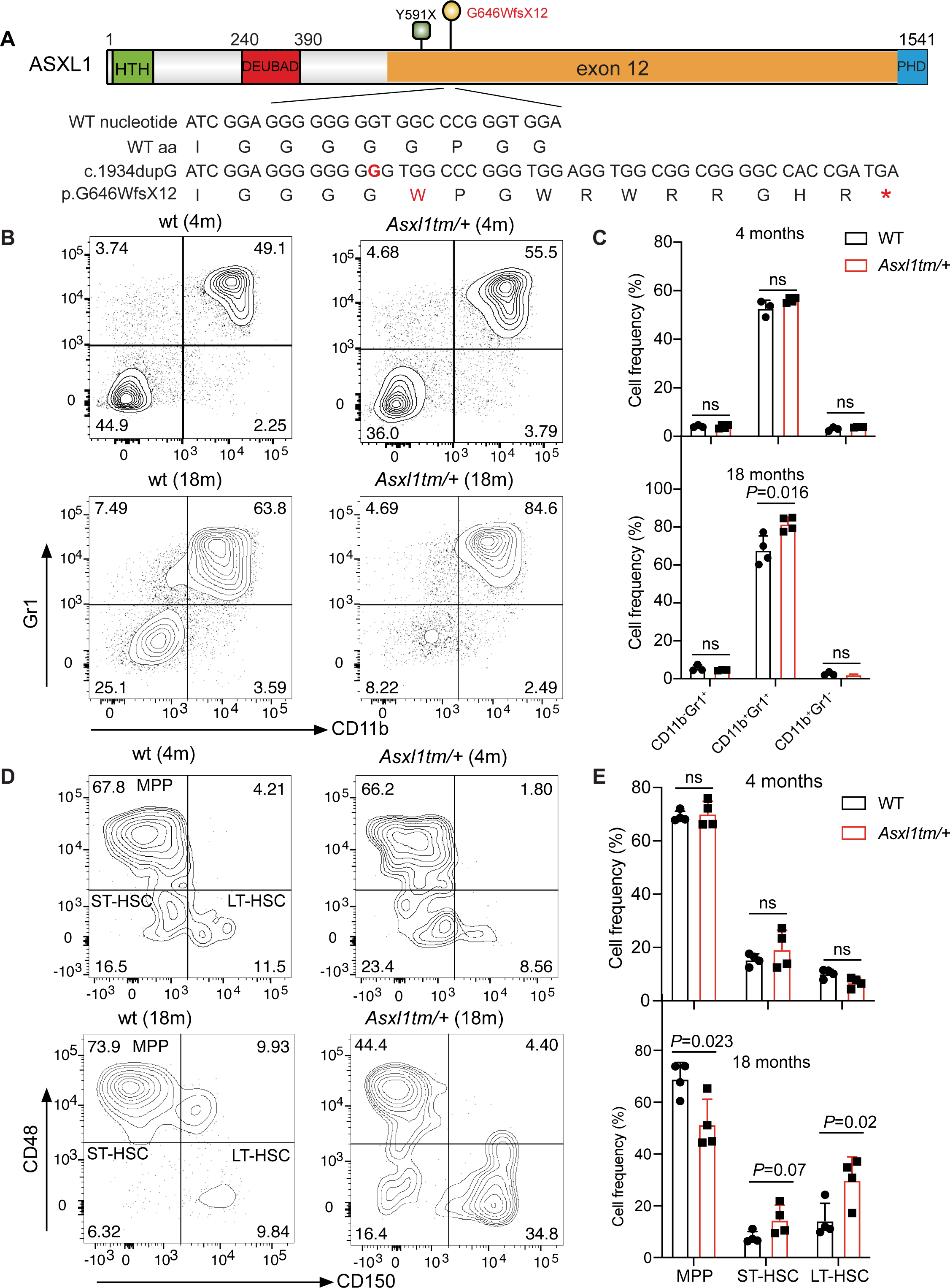
Heterozygous expression of C-terminally truncated ASXL1G646fs at physiological levels in *Asxl1tm/+* knock-in mice results in myeloid expansion with age. **A.** Schematic illustration of the mouse *Asxl1tm* knock-in mutation corresponding to the human CH/CMML-associated ASXL1G646WfsX12 mutation (ASXL1G646fs). The “G” inserted into the mutated allele is indicated in red. The protein sequence is in capitals and the asterisk represents the stop codon generated as a result of the frameshift. **B.** Flow cytometry of myeloid (Gr-1^+^/CD11b^+^) cell populations in the bone marrow of young (4 months old, *top panels*) or old mice (18 months old, *bottom panels*) WT and *Asxl1tm/+* mice. **C.** Quantification of flow cytometry in young and old WT and *Asxl1tm/+* mice (n=4 per genotype). **D.** The frequency of MPP (multipotent progenitor cells), short-term (ST) and long-term (LT) hematopoietic stem cells (HSC) in bone marrow cells of young and old mice. Lineage-negative/c-kit^+^/Sca1^+^ (LSK) cells were gated for this panel. **E.** Summary of the frequency of MPP, ST-HSCs, and LT-HSCs in bone marrow from young and old mice (n=4 per genotype). The summary graphs of flow cytometry are presented as mean ± SD. Student’s t-test is used to determine the *P* value.

Consistent with these observations, there were almost no changes in gene expression in CD11b^+^ myeloid cells from 4-month-old *Asxl1tm/+* compared to WT mice – only 2 upregulated and 2 downregulated genes based on criteria of log2 fold-change ≤1 or >1 and p-value <0.05 (***SI Appendix,* Fig S4A**). Even in older 18-month-old mice, only 40 genes were upregulated and 20 genes were downregulated in *Asxl1tm/+* compared to WT CD11b^+^ cells (***SI Appendix,* Fig S4B**). However, there were clear changes in the expression of interferon-stimulated genes (ISGs) in old *ASXL1*-mutant CD11b^+^ cells (***SI Appendix,* Fig S4B**), including *Ptgs2* (encoding prostaglandin-endoperoxide synthase 2, also known as COX2, the inducible isoform of cyclooxygenase); *Aldh1a1* and *Aldh1a7* (encoding aldehyde dehydrogenases); and *Hsp1a* and *Hsp1b* (encoding heat shock proteins) (***SI Appendix,* Fig S4B**). These findings suggest dysregulation of interferon signaling in old compared to young mutant *Asxl1tm/+* CD11b^+^ cells.

We reanalyzed RNA-seq data from long-term hematopoietic stem cells (LT-HSCs) from mice in which a different *ASXL1* mutation (ASXL1 p.E635RfsX15), was inducibly expressed from the *Rosa26* locus (24); PRJNA673672). Again, the gene expression differences compared to age-matched controls were more striking in old versus young mice, and there were marked changes in ISG expression (***SI Appendix,* Figs. S4D, S4E**). Notably, however, there was little or no overlap in either study (ours and (24)) between genes differentially expressed in young versus old cells, and the few genes whose expression was altered in common in young and old cells did not fall into any obvious functional categories (***SI Appendix*, Figs.** S4C, S4F).

Together these data indicate that unlike conventional transcription factors, mutant ASXL1 proteins may not reproducibly regulate a common set of genes. Moreover, *Asxl1* mutations seem to lead in an age-dependent manner to activation of inflammatory pathways and ISGs.

### Global decrease in H3K9me3 and H2AK119Ub in CD11b^+^ myeloid cells of old ***Asxl1tm1/+* mice**

H2AK119Ub is deposited by the RING1A and RING1B subunits of PRC1 (1, 21, 25). Since mutant ASXL1 enhances the DUB activity of BAP1 against H2AK119Ub (7), we examined the distribution of H2AK119Ub in *Asxl1tm/+* myeloid cells. Similarly, because mutant ASXL1G646fs interacts with the H3K9 methyltransferases EHMT1 and EHMT2 (**Fig. 1**) which coordinately generate H3K9me1 and me2 (8, 9), we asked whether myeloid cells from *Asxl1tm/+* knock-in mice displayed age-related changes in genome-wide distribution of the histone modification H3K9me2. We also assessed levels of the H3K9me3 modification, since H3K9me2 may serve as the substrate for deposition of H3K9me3 in heterochromatin by SUVARH1/2, SETDB1/2, and other H3K9me3 methyltransferases (26, 27); the H3K27me3 modification, since ASXL1-deficient Lin^-^c-Kit^+^ (LK) cells (which are enriched for HSPC) were reported to show decreased levels of H3K27me3 (28); and the H3K4me3 modification, since LK cells from transgenic mice expressing mutant ASXL1 proteins were reported to have decreased levels of H3K4me3 (17). We used *Drosophila* chromatin as a “spike-in” control to examine the global levels of H2AK119Ub, H3K9me2/3, H3K27me3 and H3K4me3 by CUT&RUN for in CD11b^+^ cells from old (18-to 20-month-old) *Asxl1tm/+* compared to age-matched WT mice, and divided the genome into euchromatic (Hi-C A) and heterochromatic (Hi-C B) regions by drawing on our previous Hi-C data from CD11b^+^ cells (29). H3K9me2 and H3K9me3 adorned broad regions of constitutive heterochromatin as expected (**Fig. 3A and B; *SI Appendix,* Fig. S5A**), while H3K4me3 formed sharp peaks at the promoters of transcribed genes in euchromatin (**Fig. 3C; *SI Appendix,* Fig. S5B**). H2AK119Ub and H3K27me3, histone modifications deposited by PRC1 and PRC2 respectively, are present in facultative heterochromatin, a sub-compartment of the euchromatic (Hi-C A) compartment that contains non-expressed genes that are poised to be expressed during cell differentiation or cell activation in response to external signals (**Fig. 3A and B; *SI Appendix,* Fig. S5A and B**). Both the H2AK119Ub and H3K27me3 modifications are broadly present across the gene body as well as 5’ and 3’ of genes, with the difference that H2AK119Ub marks both expressed and non-expressed genes whereas H3K27me3 primarily marks non-expressed genes (7).

**Figure 3.**
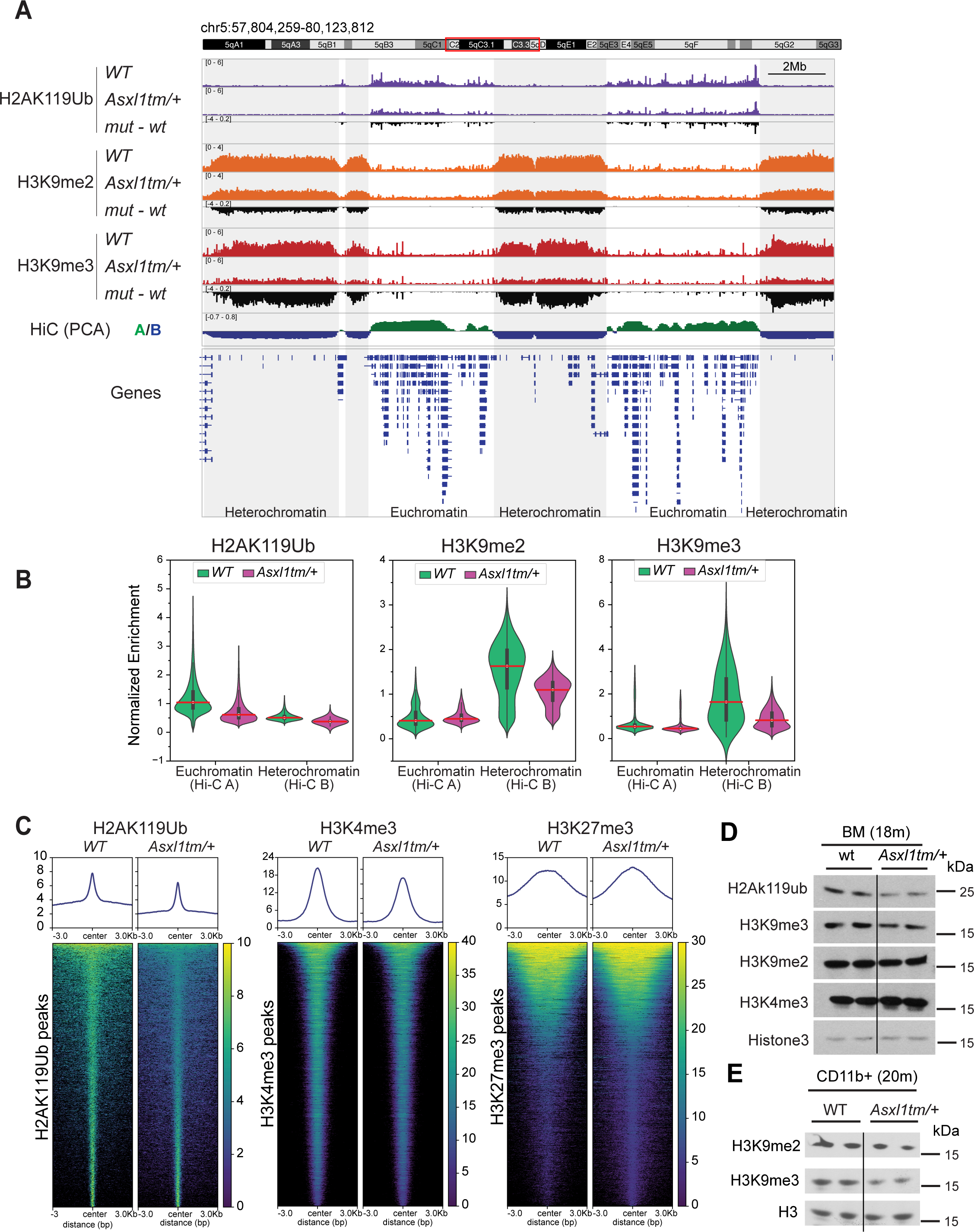
Epigenomic alterations in myeloid cells from *ASXL1tm/+* mutant mice. **A.** Genome browser views of CUT&RUN data for the indicated histone modifications in CD11b^+^ myeloid cells from aged (18- to 20-month-old) *Asxl1tm/+* mice. *Drosophila* S2 cells were used as spike-ins for normalization. A ∼26 Mb region of chr. 5 is shown. Hi-C PC1 values were used to divide the genome into euchromatic and heterochromatic compartments (positive and negative PC1 values respectively). H2AK119Ub (*tracks 1-3*) is present at peaks and broader regions in facultative heterochromatin (annotated as being in the gene-rich euchromatic (Hi-C A) compartment); H3K9me2 and me3 (*tracks 4-9*) are characteristic heterochromatic marks that cover broad gene-poor regions of the genome (*tracks 10, 11*). Compared to WT cells, *Asxl1tm/+* cells show global reduction of H2AK119Ub in euchromatic regions (*track 10, positive Hi-C PC1 values, green*), as well as global reduction of H3K9me2 and me3 in heterochromatin (*track 10, negative Hi-C PC1 values, dark blue*); this is best appreciated from the difference tracks (*3, 6 and 9*). Average IgG signals were subtracted from each sample. **B.** Violin plots showing genome-wide quantification of H2AK119ub, H3K9me2 and H3K9me3 enrichment at euchromatic and heterochromatic regions in aged *Asxl1tm/+* and wildtype mice. Signals from replicate experiments in each group were normalized to signals from *Drosophila* S2 cells that had been spiked-in to the CUT&RUN reactions; IgG signals were averaged and subtracted. The results of each sample were averaged. **C.** Heatmaps comparing the distribution of H2AK119ub, H3K4me3 and H3K27me3 signals centered at CUR&RUN peaks in CD11b^+^ cells from aged WT and *Asxl1tm/+* mice. Average IgG signals were subtracted from each sample. **D.** Western blots of acid-extracted histones from bone marrow (BM) cells from aged WT and *Asxl1tm/+* mice (18 months old) for H2AK119Ub, H3K9me2, H3K9me3 and H3K4me3 modifications. **E.** Western blots of acid-extracted histones for H3K9me2 and H3K9me3 in CD11b^+^ cells from old WT and *Asxl1tm/+* mice (20 months old). Those are from the same cells as were used for CUT&RUN. ***SI Appendix,* Fig S5C** shows a western blot and quantification of signal intensity from a similar but independent experiment with three biological replicates.

In CD11b^+^ cells of 18- to 20-month-old *Asxl1tm/+* mice compared to age-matched WT mice, we observed significant genome-wide reductions of H2AK119Ub, H3K9me2 and H3K9me3 levels (**Fig. 3A-C, *SI Appendix,* Fig. S5A**). H2AK119Ub and H3K4me3 levels were moderately reduced while H3K27me3 levels were not appreciably changed (**Fig. 3C**, ***SI Appendix,* Fig. S5B**). Western blotting of acid-extracted histones (a less sensitive technique) confirmed a perceptible reduction of the global levels of H2AK119Ub, H3K9me3 and H3K9me2 in bone marrow cells and CD11b^+^ cells of old *Asxl1tm/+* compared to age-matched WT mice (**Fig. 3D, 3E, *SI Appendix,* Fig. S5C**). In contrast, western blotting of acid-extracted histones from bone marrow cells of young (4-month-old) *Asxl1tm/+* mice showed no differences in H2AK119Ub, H3K9me2 and H3K9me3 levels compared to age-matched WT mice (***SI Appendix,* Fig. S5D**).

We also tested OCI-AML2 cells transduced with full-length wildtype or mutant (mt) ASXL1 Y591X, and with GFP as a control. OCI-AML2 cells have no baseline mutations in *ASXL1* or *TET2*, but western blotting of acid-extracted histones from the transduced cells showed a substantial reduction of both H3K9me3 and H2AK119Ub, with no obvious reduction in H3K9me2; both effects were intensified when BAP1 was co-expressed (***SI Appendix,* Fig. S6A**). Consistent with our previous observation in HEK293T cells, the ASXL1 mutant protein is highly expressed and much more stable than the full-length proteins in all cell types tested: OCI-AML cells, K562 cells and HEK293T cells (***SI Appendix,* Fig. S6B-D**).

Taken together, these data show that the global reduction in H2AK119Ub, H3K9me2 and H3K9me3 levels occurs progressively with age, in parallel with the observed increases in myeloid cell frequencies, the dysregulation of other hematopoietic lineages and the alterations in differential gene expression with age (**Fig. 2, *SI Appendix,* Fig. S3, S4**).

### Old *Asxl1tm/+* mice display increased expression of transposable and repetitive elements, increased levels of R-loops and DNA damage

Decreased H2AK119Ub and H3K9me2/3 levels in heterochromatin are associated with increased expression of transposable elements (TEs) and satellite repeats that are normally tightly suppressed in healthy cells (21, 25–27). We compared TE expression in CD11b^+^ cells of young (4-month) and older (20-month) *Asxl1tm1/+* vs WT mice by qRT-PCR, with DNase I treatment to eliminate DNA contamination. In CD11b^+^ cells from 18-month-old but not 4-month-old *Asxl1tm/+* mice compared to age-matched controls, we observed a significant increase in the expression of major and minor satellite repeats and LINE-1s (L1s) by qRT-PCR (**Fig. 4A**). RNA-seq of total RNA from CD11b^+^ cells of old *Asxl1tm1/+* mice revealed increased expression of an L1 subfamily (L1MEi) as well as the internal (int) and LTR regions respectively of two subfamilies of endogenous retroviruses (RMER17C-int and IAPEY4 LTR)) (**Fig. 4B**). In each case, increased TE expression was even more pronounced in LK cells of *Asxl1tm/+* mice, compared to LK cells from aged-matched wildtype control mice (**Fig. 4C and D**). Taken together with ***SI Appendix,* Fig. S4**, the data suggest that ASXL1 mutations, especially in HSPC, intensify the increased TE expression and inflammation associated with normal aging.

**Figure 4.**
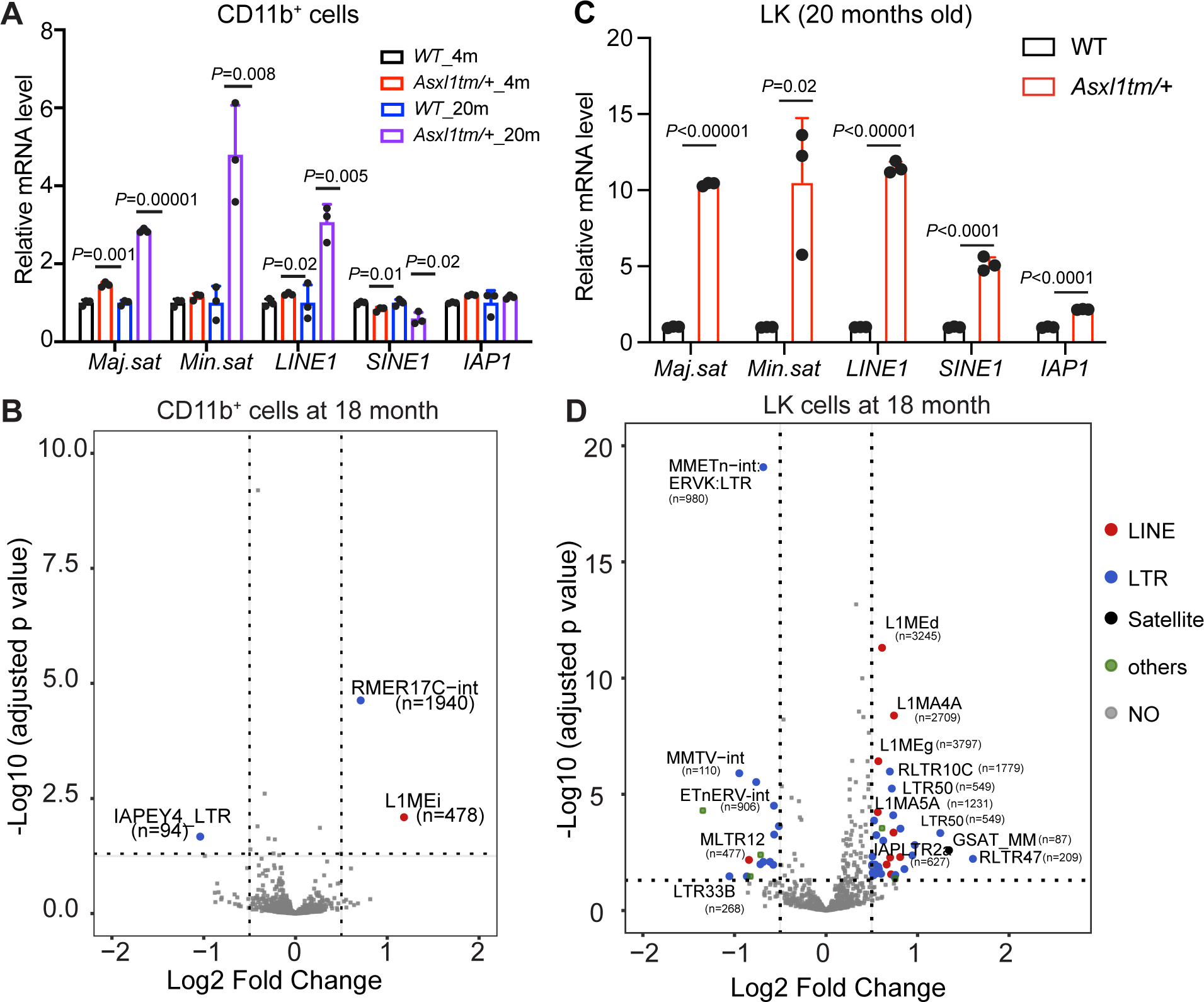
Increased expression of transposable elements and satellite repeats in aged *ASXL1tm/+* mutant cells. **A.** RT-qPCR of repetitive elements in CD11b^+^ cells from young and old *Asxl1* mice. Total RNA isolated from CD11b^+^ cells from young (4 months old) and old (18 months old) *Asxl1 tm/+* and WT mice was treated with DNase I and expression of satellite repeats and TEs was assessed by quantitative RT-PCR (qRT-PCR). **B.** Volcano plot of RNA-seq data showing limited changes of LINE-1s and LTRs in CD11b^+^ cells from old (18 months old) *Asxl1tm/+* mice compared to CD11b^+^ cells from age-matched WT mice. **C.** Quantitative RT-PCR of repetitive elements in LK (Lineage^-^c-Kit^+^) cells from 20 months-old mice. Comparison with the data from CD11b^+^ cells (**A**) shows a greater increase in expression of TEs and major and minor satellite repeats in LK cells from old *Asxl1 tm/+* mice compared to LK cells from age-matched WT mice. **D.** Volcano plot of RNA-seq data showing markedly increased expression of LINE-1s, LTRs and GSAT_Mm satellite repeats in LK cells from old (18 months old) *Asxl1 tm/+* mice compared to age-matched WT mice. The increase is considerably greater than that shown in (**B**) for CD11b^+^ cells from old *Asxl1 tm/+* mice. P<0.05, Log2FC+/-0.5. Student’s t-test is used to determine the *P* value.

Cells with decreased H2K119Ub or decreased DNA or H3K9 methylation often display increased levels of R-loops and DNA damage (11, 26, 27, 30). In *C. elegans* lacking the SETDB1 and SUV39H homologs MET-2 and SET-25, R-loops were substantially enriched over repeat elements that were derepressed in the absence of H3K9me3 (27), while in humans and other species, a variety of genomic repeats have been reported to be enriched for R-loops (31, 32). Using recombinant V5-tagged catalytically inactive (D210N) RNASE H1 protein (dRNase H1) in flow cytometry of permeabilized cells, we observed that R-loops levels increased significantly in CD11b^+^ myeloid cells from young (4-month-old) and more strikingly in old (20-month-old) *Asxl1tm/+* compared to WT mice (***SI Appendix,* Fig. S7A**).

Increased R-loops are highly correlated with DNA damage (32, 33); specifically, excess R-loops are excised by the xeroderma pigmentosa proteins XPG and XPF, which can generate both single-strand and double-strand DNA breaks (33). However, a significant increase in double-stranded DNA breaks, assessed by flow cytometry for γH2AX, was observed only in old *Asxl1tm/+* compared to WT mice (***SI Appendix,* Fig. S7B**). While the correlation of increased TE and satellite repeat expression with increased R-loops and DNA damage is intriguing, long-reads sequencing and other emerging technologies will be needed to ask if the genomic location of DNA damage associated with newly emerging R-loops can be connected directly to increased expression of specific uniquely mapped TEs and satellite repeats.

### CMML patients show increased expression of interferon-stimulated genes (ISGs) and TEs located in heterochromatin

*ASXL1* mutations that result in C-terminally truncated proteins are associated with age-related CH, as well as with CMML, MDS and AML (12, 13). In CMML, approximately 40% of patients have truncating *ASXL1* mutations and these mutations are associated with a proliferative phenotype, resistance to epigenetic therapies and shortened overall survival (<2 years) (34). We reanalyzed published RNA-seq data (34) from bone marrow mononuclear cells (primarily malignant monocytes) of *ASXL1*-mutant CMML patients (**Fig. 5**); the variant allele frequencies of the *ASXL1* mutations (***SI Appendix,* Fig. S1**) ranged from 31-48%.

**Figure 5.**
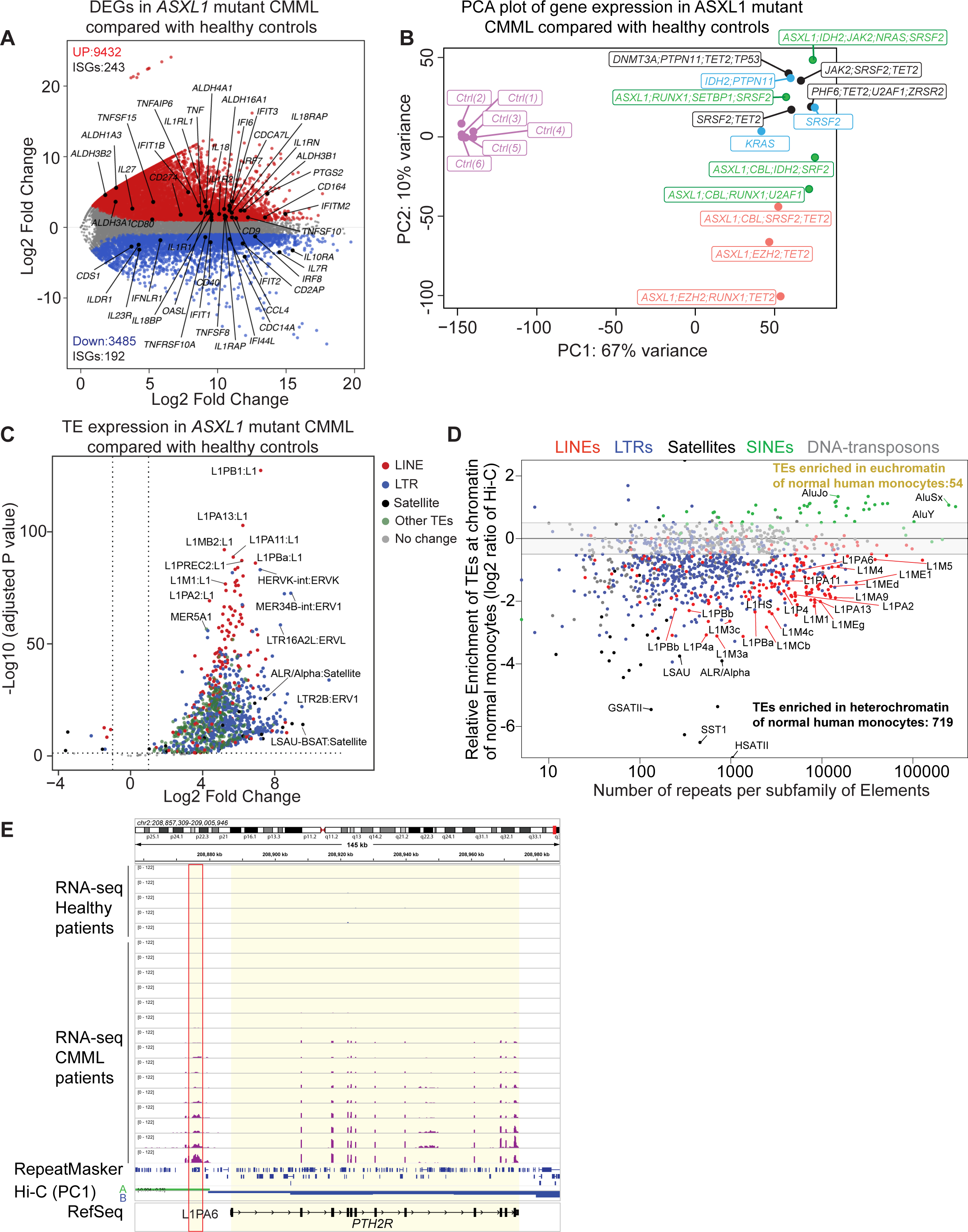
Increased expression of Interferon-Stimulated Genes (ISGs) and transposable elements in monocytes from ASXL1-mutant CMML patients. **A.** Differential gene expression in cells from ASXL1-mutant CMML patients. The MA-plot highlights differentially expressed genes (DEGs; red dots, expressed at high levels in ASXL1-mutant CMML; blue dots, genes downregulated in ASXL1-mutant CMML compared to healthy controls). Differentially expressed ISGs are shown as black dots. The numbers of DEGs and differentially expressed ISGs are indicated. **B.** Principal Component Analysis (PCA) plot of RNA-seq in monocytes from healthy controls versus ASXL1-mutant CMML patients bearing different mutations in addition to ASXL1. **C.** Volcano plot of differentially expressed transposable elements (TEs) in monocytes from ASXL1-mutant CMML patients compared to healthy controls. The bulk expression of TE families was analyzed from Poly (A) RNA-seq data using the TE transcripts package. Each dot represents a specific TE-subfamily. Color codes show the different classes of TEs. Differentially expressed TEs were considered as log2 fold change (log2FC) of +/- 1 and a padj value <0.05 (calculated by adjusting the p-value with the false discovery rate [FDR]). **D.** Distribution of TE subfamilies in euchromatin or heterochromatin in normal human monocytes. For each subfamily of TEs (represented as a single dot), we counted the number of individual TE copies located in euchromatin (positive Hi-C PC1 values) or heterochromatin (negative Hi-C PC1 values). The log2 ratio of the number of TE copies is plotted on the y-axis, while the total number of copies found in the genome (T2T) is shown on the x-axis. Enriched TEs were defined as log2 ratio > or < +/- 0.5. **E.** Genome browser view showing that a uniquely mapped *L1PA6* element in euchromatin (red rectangle) is variably upregulated in monocytes from a subset (10/15) *ASXL1*-mutant CMML patients; a downstream gene, *PTH2R,* located in heterochromatin in normal human monocytes (dark blue), also displayed higher expression in *ASXL1*-mutant cells, and the extent of *L1PA6* expression correlated closely with the extent of *PTH2R* upregulation.

Thousands of genes were up-regulated and downregulated in samples from ASXL1-mutant CMML patients compared to healthy controls (**Fig. 5A**; ***SI Appendix,* Fig. S8A, S8B**); many of these were interferon-induced genes (ISGs), consistent with the high levels of inflammation characteristic of CMML (12) (**Fig. 5A**). A PCA plot showed that the gene expression patterns of healthy control individuals clustered tightly together but were separated in the PC1 dimension from the gene expression patterns of CMML patients harboring ASXL1 mutations (**Fig. 5B**).

We also analyzed changes in TE expression in *ASXL1*-mutant CMML patients (**Fig. 5C**). Compared to healthy individuals (controls), *ASXL1*-mutant CMML patients showed a striking increase in the expression of many TE subfamilies including L1s and LTRs (human endogenous retroviruses, HERVs) as well as alpha (ALR-ASAT) and beta (LSAU) satellite repeats and DNA transposons (**Fig. 5C**; ***SI Appendix,* Fig. S8C-E**). Notably, these TE subfamilies and repeat classes are located primarily in heterochromatin (**Fig. 5D**), when plotted using available Hi-C data from normal human monocytes (35). The data suggest that with more extensive analyses involving a broader spectrum of patients, it might be possible to use RNA-seq and repeat/TE expression to subclassify CMML for disease stratification or for prognostic or therapeutic purposes. HERVK LTRs were prominent among the upregulated TEs and their relevance to cancer immunotherapy has been noted previously: specifically, HERVK-int expression is potentially immunogenic (36), and expression of HERVK loci has been used to stratify lung cancer patients (37).

Since TEs can act as gene promoters or enhancers (38, 39), we asked if the increased TE expression observed in CMML patients was accompanied by changes in the expression of nearby genes and TEs. Only a very few individual upregulated TEs could be mapped uniquely using the available RNA-seq data (50x50 paired-end reads) (34); all of these were located in heterochromatin in healthy human monocytes and massively but variably upregulated in CMML (***SI Appendix,* Fig. S8C-E**). One of these uniquely-mapped elements – an evolutionarily young L1PA6 element (40) – was located 5’ of the *PTH2R* gene, and its increased expression correlated with increased *cis*-expression of the *PTH2R* gene in *ASXL1*-mutant CMML (**Fig. 5E**). The clinical significance of increased *PTH2R* expression is not known, but the data are reminiscent of the heterochromatin-to-euchromatin switch that we previously documented in expanded myeloid cells of inducible *Tet1/2/3*-deficient mice (29).

Since CMML is a highly inflammatory disease, with 30% of patients showing concomitant systemic inflammatory manifestations (12), we assessed inflammatory phenotypes by examining interferon-stimulated genes (ISGs), which include genes encoding IRF family members, pro-inflammatory cytokines (IL6, IL1β) and cytokine and chemokine receptors. Indeed, as in older myeloid cells from *Asxl1tm/+* mice (***SI Appendix,* Fig. S4**), a substantial proportion of differentially expressed genes in CMML patients were ISGs (**Fig. 5A**). Notable among these were *PTGS2* (encoding COX2, the inducible cyclooxygenase isoform) and genes belonging to the *ALDH* aldehyde dehydrogenase family, including *ALDH1A3* and *ALDH16A1* (**Fig. 6A, B**). As noted above, *Ptgs2* and related *Aldh*-family genes were also found upregulated in myeloid cells from old *Asxl1tm/+* mice (***SI Appendix,* Fig. S4B, Fig. 6C**). Both *Aldh1a1* and *Aldh1a7* genes are located in heterochromatin (Hi-C B compartment) in normal myeloid cells and are upregulated together in CMML (**Fig. 6D**), and low ALDH levels are associated with more favorable outcomes in AML (41).

**Figure 6.**
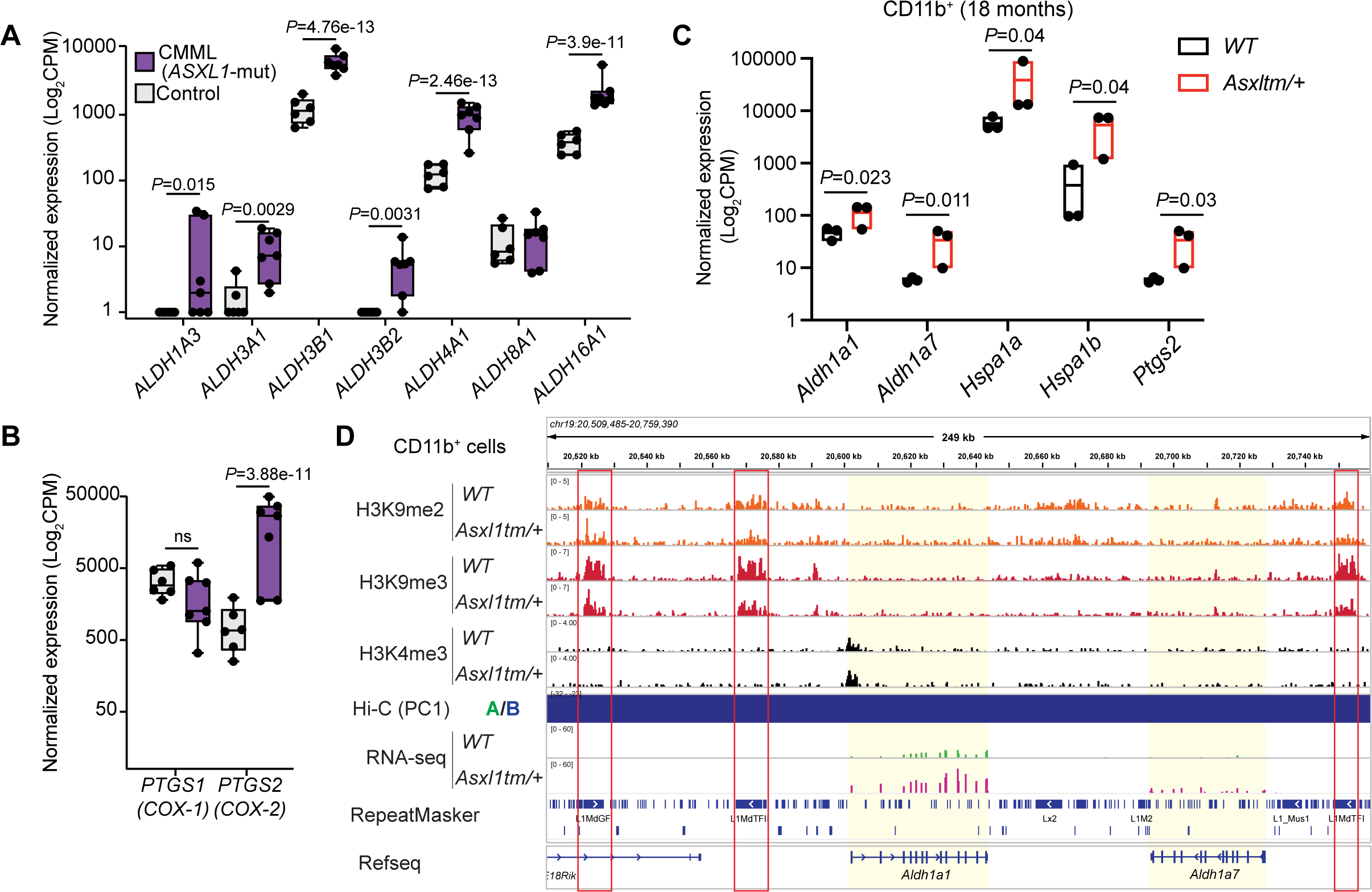
Increased expression of *ALDH*-family genes in monocytes from *ASXL1*-mutant CMML and heterochromatin-localized *Aldh1a* genes in myeloid cells from *Asxl1tm/+* mice. **A.** Normalized expression of *ALDH*-family genes in *ASXL1*-mutant CMML. Data are presented as Log_2_CPM (counts per million reads) after normalization from RNA-seq. **B.** Normalized expression of *PTGS1* and *PTGS2* in *ASXL1*-mutant CMML. **C.** Normalized expression of ALDH-family genes upregulated in old (18 months) *Asxl1tm/+* CD11b^+^ cells (*this study*). Data are shown as Log_2_CPM (counts per million reads) after normalization from RNA-seq data. **D.** Genome browser view showing that *Aldh1a1, Aldh1a7* and three adjacent L1s are buried in heterochromatin (defined by Hi-C PC1 values) in *WT* mouse CD11b^+^ cells. The entire region shows a perceptible decrease in levels of H3K9me2/me3 (*top tracks*), and *Aldh1a1* expression (and H3K4me3 levels at the *Aldh1a1* promoter) are upregulated in myeloid cells from *Asxl1tm/+* mice (RNA-seq tracks).

*ALDH*-family and *PTGS2* genes were prominent among the ISGs upregulated both in our mouse model of CH in older *Asxl1tm/+* mice and in monocytes from CMML patients (***SI Appendix,* Fig. S4B**; **Fig. 6A-C**). Several ALDH-family genes show high expression in AML, and low *ALDH1A1* expression correlates with improved survival in AML(41, 42). Mouse LSK cells express high levels of *Aldh1a1* mRNA and ALDH1A1 protein compared to total bone marrow cells, and *Aldh1a1* and *Aldh3a1* together regulate hematopoietic (particularly B cell) lineage specification in mice (43). However, following HSPC differentiation into mouse CD11b^+^ cells or differentiated human monocytes, these genes become localized to heterochromatin, which contains the majority of L1s and LTRs (**Fig. 6D**).

Together these findings suggest that ASXL1 is not a conventional transcriptional regulator that directly controls gene expression by binding to DNA or acting as a coactivator/ corepressor for DNA-binding transcription factors. Rather, the decrease in histone modifications in facultative and constitutive heterochromatin may drive stochastic increases in gene and TE expression. On occasion (as for *PTH2R* in CMML), we observe parallel increases in the expression of TEs and nearby genes in *cis*, suggesting a connection between TE and *cis*-expression of 3’ genes that is yet to be defined. This hypothesis may explain why reproducible changes in gene expression are rarely observed in *ASXL1*-mutant CMML (34) or in different mouse models of ASXL1 mutations (***SI Appendix,* Fig. S4**).

## Discussion

Here, we show that CH/CMML-associated ASXL1 mutant proteins co-immunoprecipitate (co-IP) not only with the known interactors BAP1, HCF1, OGT and FOXK1/2, but also with a set of heterochromatin-related proteins that were not previously described as interacting with ASXL1. Notable among these are EHMT1 and EHMT2, a dimer of H3K9 methyltransferases that coordinately generate H3K9me2 (8, 9). EHMT2 co-IPs with both ASXL1 and BAP1, implying the existence of an ASXL1-BAP1-EHMT1-EHMT2 complex that may contain either EHMT1 and EHMT2 homodimers or the EHMT1-EHMT2 heterodimer (**Fig. 1H**, ***SI Appendix,* Fig. S9A**). The inability of BAP1 to co-IP with OGT, HCF1, FOXK1 or FOXK2 implies the existence of a distinct ASXL1-BAP1-OGT-HCF1 complex (**Fig. 1H**, ***SI Appendix,* Fig. S9A**) that may also contain FOXK1 and FOXK2 (15, 44).

The ASXL1-BAP1 complexes appear to be involved in regulating numerous histone modifications that are dysregulated in cells bearing ASXL1 truncation mutations associated with CH, CMML and AML. Myeloid cells from old *Asxl1tm/+* mice showed a genome-wide decrease in mono-ubiquitylated H2AK119Ub, consistent with the well-established ability of C-terminally truncated ASXL1/2/3 proteins to potentiate BAP1 deubiquitinase activity (7) through the interaction of ASXL DEUBAD domains with the UCH-CC1 domains of BAP1 (44–46). HCF1 is a tight partner of OGT that is part of the MLL H3K4 methyltransferase complex (47), and the moderate reduction of H3K4me3 in myeloid cells from old *Asxl1tm/+* mice (**Fig. 3**) presumably reflects the presence of mutant ASXL1 in the canonical ASXL1-BAP1-OGT-HCF1 complex (***SI Appendix,* Fig. S5**). Finally, the genome-wide decrease in H3K9me2 and me3 levels in myeloid cells of old *Asxl1tm/+* mice – a model of age-related CH is likely to be mediated through the ASXL1-BAP1-EHMT1/EHMT2 complex although the mechanisms remain to be defined. Cryo-EM studies show that BAP1 tethers the ASXL1 DEUBAD domain to nucleosomes through its interaction with the nucleosome acidic patch (45, 46), suggesting that the decreases in H2AK119, H3K9 and H3K4 modification mediated by mutant ASXL1 all require the ASXL1-BAP1 interaction.

Mutant ASXL1 also interacted with three other well-established heterochromatin-associated proteins, NPM1 (nucleophosmin), HP1γ and CENPV. The nucleolar protein NPM1 is localized to heterochromatin and recurrently mutated in AML; mutant NPM1 loses its C-terminal nucleolar localization signal and becomes localized to the cytoplasm (23). HP1γ, also a heterochromatin-associated protein that binds H3K9me3, was listed among the interaction partners of ASXL1 E635RfsX15 and ASXL1 Y588X in previous studies but was not further pursued (17, 18). The role of the ASXL1-CENPV (centromere protein V) interaction is less clear, primarily because the functions of CENPV have not been extensively investigated. In HeLa cells, overexpression of CENPV leads to hypercondensation of pericentromeric heterochromatin and altered H3K9me3 distribution, whereas CENPV depletion correlates with a number of chromosomal abnormalities in metaphase (48).

EHMT1 and EHMT2 primarily regulate H3K9me1/2 in mice, with only a minor effect on H3K9me3 (9, 49, 50). Notably, however, the decrease in H3K9me3 that we observed in our study was more prominent than the decrease in H3K9me2, suggesting the involvement of H3K9me3 methyltransferases such as SUVARH1/2 and/or SETDB1/2. It will be important in future studies to define the structures and activities of the two postulated complexes containing mutant ASXL1; determine whether mutant ASXL1 in the ASXL1-BAP1-EHMT1-EHMT2 complex inhibits EHMT1/EHMT2 H3K9me1/me2 activity; and identify the H3K9me3 methyltransferases whose reduced activity is responsible for the observed reduction in H3K9me3 levels in *Asxl1tm/+* cells.

Our data prompt the unifying hypothesis that all three of the most frequent mutations in CH and hematopoietic malignancies – in *DNMT3A*, *TET2* and *ASXL1* – involve a progressive loss of heterochromatin integrity (21) (***SI Appendix,* Fig. S9B**). TET deficiency in a variety of different cell types is associated with decreased DNA methylation in heterochromatin (11, 51); likewise, DNMT3A deficiency results in decreased DNA methylation genome-wide and *Dnmt3a* and *Tet2* deficiency cooperate in mice to induce more rapid dysregulation of myeloid phenotypes and reduced survival (11). Moreover, reduction of DNA methylation or H3K9me2/me3 modifications in constitutive heterochromatin is associated with a variety of cellular and genomic aberrations – increased expression of TEs and other repetitive elements, increased R-loops, increased mutations and genome instability, and increased expression of genes related to inflammation (11, 21, 26, 27, 51). Double *Tet2, Tet3*-deficient (*Tet2/3 DKO*) iNKT cells (11), *Tet2/3 DKO* B cells (32) and ASXL1-mutant myeloid cells (*this study*) display increased expression of TEs and satellite repeats, increased R-loops and increased DNA damage. *Tet iTKO* mESC display chromosome missegregation and aneuploidies (52); and *Tet2-/-* peritoneal macrophages (53), *Tet2-/-* bone-marrow-derived macrophages (BMDM) (54), *Tet iTKO* BMDM (55) and *DNMT3A*- or *TET2*-deficient human monocyte-derived macrophages (56) all show increased expression of pro-inflammatory cytokines, cytokine receptors and interferon-stimulated genes (ISGs). These changes are manifestations of heterochromatin “weakening” or “dysfunction” (11, 21, 51).

We postulate that during age-dependent CH evolution and CH to CMML progression, increased expression of TEs and heterochromatin-associated genes occurs through slow, stochastic release from heterochromatin-mediated silencing, followed by positive selection for clones with increased expression of specific protein-coding genes and TEs in heterochromatin. Among these are *ALDH*-family genes; notably, high *ALDH* levels in AML are associated with poor patient survival (41, 42), suggesting that greater derepression of heterochromatic genes (including *ALDH1A* genes in CMML and *Aldh1a* genes in *Asxl1tm/+* mice) and TEs in both aging mice with a CH-associated *Asxl1* mutation and humans with CMML may worsen outcomes in age-related CH and an aging-related cancer that emerges from CH. Likewise, we previously documented the expansion of new *Tet iTKO* CD11b^+^ myeloid populations expressing higher levels of clustered *stefin/cystatin* genes that were originally localised to heterochromatin but became involved in a limited heterochromatin-to-euchromatin switch (29). In this case as well, high expression of cystatin B (CSTB) in AML patients correlated with poor survival (29). Our findings suggest that a higher degree of heterochromatin weakening or dysfunction, leading to stochastic re-expression of genes otherwise suppressed in heterochromatin, may be associated with adverse patient outcomes in AML and potentially in other cancers as well, and that analysis of RNA-seq data for increased TE expression may be warranted as a prognostic biomarker for many cancers and inflammatory diseases.

DNA methyltransferase inhibitors (DNMTi) such as azacitidine, decitabine and decitabine with the cytidine deaminase inhibitor cedazuridine are US FDA approved therapies for myeloid neoplasms such as MDS, CMML and AML (57). The clinical improvement observed after treatment of responder patients with DNMTi does not appear to be due to preferential elimination of mutant HSPC clones (e.g. by the immune system), alteration in mutational allele burdens, or a decrease in the differentiation capability of mutant HSPC (58); rather, it is thought that these agents likely act in the short term by increasing inflammation through “viral mimicry”, thereby potentiating anti-tumor responses (59). Our data suggest that in the longer-term, therapy with DNMTI would not improve outcomes for patients with MDS, AML or CMML: the slow, genome-wide, replication-dependent reduction of DNA methylation that accompanies such therapies might worsen heterochromatin dysfunction and disease progression. Rather, it would be ideal, if possible, to devise therapies that delay the slow loss of DNA and H3K9 methylation during the progression of CH to hematopoietic malignancies.

## Materials and Methods

Detailed materials and methods are described in *SI Appendix*, *Materials* and *Methods*.

### Mice

The *Asxl1tm* strain of mutant knock-in mice were generated in the laboratory of Dr. Hwei-Fang Tien. The cognate mouse mutation is predicted to be c.1925dupG; p.G643WfsX12, encoding 654 amino acids mimicking the most common form of human mutant ASXL1 protein p.G646WfsX12. Mice were housed in a pathogen-free animal facility at the La Jolla Institute. They were used according to protocols approved by the Institutional Animal Care and Use Committee (IACUC).

## Data and Software Availability

Next-generation sequencing have been deposited in the Gene Expression Omnibus (GEO) repository, accession of RNA-seq in CD11b^+^ cells and LK cells from mice is GSE274878, the RNA-seq of CMML patients is GSE274876, and the CUT&RUN data is GSE274875.

## Acknowledgments

We thank Steve Henikoff and Kami Ahmad (Fred Hutchinson) for reagents, protocols, suggestions and support regarding CUT&RUN experiments, and the Department of Laboratory Animal Care (DLAC) and the animal facility for excellent support. We also thank C. Kim, S Alarcon and Z. Mikulski and colleagues of the La Jolla Institute (LJI) Flow Cytometry, Next-Generation Sequencing and Microscopy Core Facilities for help with cell sorting, next-generation sequencing, and microscopy respectively. The FACSAria II Cell Sorter (S10 RR027366), the CyTOF Mass Cytometer (S10 RR027366), the NovaSeq 6000 (S10OD025052) and the HiSeq 2500 (S10 OD018499) were acquired through the Shared Instrumentation Grant (SIG) Program. This research was supported by NIH grants R35 CA210043 to A.R., R01 CA272496 to M.P., NIH NIGMS R35GM147554 to S.A.M., NHMRC Investigator Grant (GNT1173711) and the Mater Foundation to G.J.F., NSC 112-2314-B-075A-004- and 112-2314-B-002-095-MY3 to W.-C.-C, and MOHW109-TDU-B-211-134009 to W-C.C. and H.-F. T. H.S. was supported by the Pew Latin-American Fellows Program from The Pew Charitable Trusts, and by a Fellowship from the California Institute for Regenerative Medicine.

## Author Contributions

Z.D. designed and performed experiments, analyzed data and prepared the sequencing libraries; H.S. and L.J.A. performed computational analyses of the RNA-seq and CUT&RUN data; C.B. assisted with cell isolation and flow cytometry; W-C.C. and H.F.T. provided the ASXL1 mutant knock-in mouse strain; G.J.F. advised on and interpreted the data on transposable element expression together with H.S., S.A.M. performed the mass spectrometry experiments and analyzed the data, J.F., M.B. and M.P. provided RNA-sequencing data for CMML samples and healthy controls. Z.D., H.S. and A.R. wrote the manuscript, with all authors contributing to writing, editing and providing feedback on the manuscript.

## Classification

Research Article

## Supporting Information

### Materials and Methods

#### Plasmids

Mammalian cytomegalovirus (CMV) promoter (pCMV)-driven plasmids that encode full-length human ASXL1 and BAP1, C-terminally FLAG-tagged ASXL1 p.G646Wfs*12 (Addgene 74245) and p.Y591X (Addgene 74261) could be obtained from Addgene. The MSCV-T2A-Puromycin vectors encoding 3xFLAG-tagged full-length ASXL1 and truncated ASXL1 were generated by PCR amplification followed by sub-cloning using Gibson assembly.

#### Flow cytometry

Antibody staining and flow cytometric analysis was performed as previously described (1). Briefly, bone marrow cells were flushed out of femurs and tibiae in Hanks balanced salt solution (HBSS) containing 2% FBS and 10M HEPES. Red blood cells were depleted prior to flow cytometry. Cells were stained with monoclonal antibodies in FACS buffer (PBS containing 3% heat-inactivated FBS, 2 mM EDTA and 0.1% (w/v) sodium azide. All antibodies were purchased from Biolegend and BD Bioscience. The following monoclonal antibodies were used: PerCP/Cy5.5 anti-mouse Cd11b, BV650 anti mouse Gr1, APC anti-mouse CD3, BV421 anti-mouse B220. As a lineage cocktail, biotinylated antibodies against CD3ε, CD45R/B220, Mac-1/CD11b, Gr-1, Ter-119 (all from Mouse hematopoietic Lineage Biotin Panel, Biolegend) and IL-7Rα were used in combination with a Streptavidin-PerCP-Cy5.5 conjugate (Biolegend). Live/Dead eFluor 780 (Invitrogen) was used to exclude dead cells. Flow cytometric analyses were performed using FACS Celesta (BD Biosciences) and data were analyzed using FlowJo software. Cd11b^+^ (Mac-1^+^) cells were isolated from unfractionated bone marrow using CD11b MicroBeads (Miltenyi Biotec) following the manufacturers’ instructions.

#### Immunoprecipitation and mass spectrometry

For sample generation, nuclear extracts were prepared from approximately 70 million HEK293T cells transfected with untagged BAP1 and mutant ASXL1G646fs bearing a C-terminal 3xFLAG-tag as previous description (2). The cells were washed twice with cold PBS, and then resuspended in hypotonic buffer (10 mM HEPES, 1.5 mM MgCl_2_, 10 mM KCl). After 15 min incubation on ice, cells were pelleted (650*g*, 5 min), and resuspended in isotonic buffer (10 mM Tris pH 7.5, 2 mM MgCl_2_, 3 mM CaCl_2_, 0.3 M sucrose). Cells were disrupted using a Dounce homogenizer, followed by centrifugation (10,000*g*, 20 min) to collect pellets containing nuclei. Pellets were resuspended in nuclear lysis buffer (10 mM HEPES pH 7.9, 400 mM NaCl, 1mM EDTA and 0.5mM DTT) supplemented with benzonase (500 U/ml), and incubated at 4°C for 30 min on a rotator. Debris was removed by centrifugation at 13,000 rpm for 15 min, and the supernatant (nuclear fraction) was recovered. An equal volume of 2x conversion buffer (10 mM Tris-HCl pH 7.5, 280 mM NaCl, 1 mM EDTA, 1 mM EGTA, 0.2% Triton X-100, 10% glycerol) was added to the nuclear proteins. Anti-FLAG M2 magnetic beads (Sigma M8823) were washed with RIPA buffer before adding 1 µl beads/ mg protein to protein samples with benzonase and inverting continuously at 4°C overnight. Reactions were carried out overnight at 4°C and washed three times with wash buffer I (50 mM Tris HCl, pH 7.4, with 150 mM NaCl, 1 mM EDTA, and 1% TRITON-X-100) and wash buffer II (50 mM Tris-HCl pH 7.4, 150 mM NaCl). 5% of beads suspension was aliquoted and heated at 90°C in 1× LDS sample buffers with 5% 2-mercaptoethanol to elute proteins. Immunoprecipitated proteins were analyzed using immunoblotting and silver staining. The remaining protein-beads complexes were used for LC-MS/MS as previously described (3). Protein quantification was done using Pierce™ BCA Protein Assay Kits (Thermo Fisher Scientific, 23225). Proteins were processed and analyzed as previously described with minor modifications (3). Samples were analyzed on an Orbitrap Eclipse mass spectrometer (Thermo Fisher Scientific).

#### Mass Spectrometry data analysis

Raw data were processed using MaxQuant software (4) using the default settings and searched against the human UniProt database (12/2017) with common contaminant entries. The settings used for MaxQuant analysis were: enzymes set as LysC/P and Trypsin/P, with maximum of 2 missed cleavages; fixed modification was Carbamidomethyl (Cys); variable modifications were Acetyl (Protein N-term) and Oxidation (Met); label-free quantification was used with a minimum ratio count of 2 and classic normalization; False Discovery Rate for both protein and peptide identification was 0.01. The ‘Match between runs” feature was enabled. The proteinGroups file from the MaxQuant analysis was used as the input for all downstream analyses. Gene names that were not automatically assigned by MaxQuant were manually added from their Entrez protein IDs. Each of the two replicates within a condition were annotated. LFQ intensities were log2 transformed and proteins with less than 2 LFQ intensity values across both conditions were filtered out. Missing values were imputed with the minimum detected intensity. A two-tailed t-test with a p-value cutoff of 0.05 and FDR of 0.01 was conducted alongside the calculation of log2 fold changes for construction of the volcano plot.

#### Immunostaining

HEK293T cells were seeded on round coverslips treated with L-poly-lysine in 24-well plate and transfected with plasmids. Cells were fixed with 4% formaldehyde for 15 min at room temperature and blocked with 3% normal goat serum and in PBST (0.2% Triton-X 100 in 1X PBS) for 20 min at room temperature. Staining was carried out by incubating overnight with primary antibodies (1: 500) in PBST supplemented with 3% normal goat serum. The cells were rinsed three times with 1X PBS and incubated with fluorochrome-conjugated secondary antibodies (1:1,000) in the blocking buffer for 1 h at room temperature. The cells were rinsed three times with 1X PBS and coated with ProLong Gold antifade mounting medium with DAPI (4’,6-diamidino-2-phenylindole, 100 ng/ml) (Life Technologies, P-36931) on slides. Images were obtained on an LSM 880 laser-scanning confocal microscope.

#### RNA extraction and qRT-PCR

Total RNA was isolated with RNeasy Plus Mini Kit or with Trizol following the manufacturers’ instructions. To prevent contamination from genomic DNA when using the Trizol, the purified RNA was subjected to DNase I treatment (1 U/10 ng RNA) for 30 minutes at 37°C, then the DNase I was inactivated by adding 5 mM EDTA and heating at 75°C for 10 minutes. RNA was concentrated using 5 μg linear acrylamide and 0.5 M ammonium acetate, followed by 100% ETOH to precipitate RNA by centrifugation at 14,000 rpm for 15 min at 4°C. cDNA was synthesized using SuperScript III reverse transcriptase, and qRT-PCR was performed using SYBR Green Master Mix (Applied Biosystems A25741) on a StepOnePlus Real-time PCR system (Thermo Fisher Scientific). Gene expression was normalized to 18s rRNA. Primers for qRT-PCR are listed below as previous description (5).

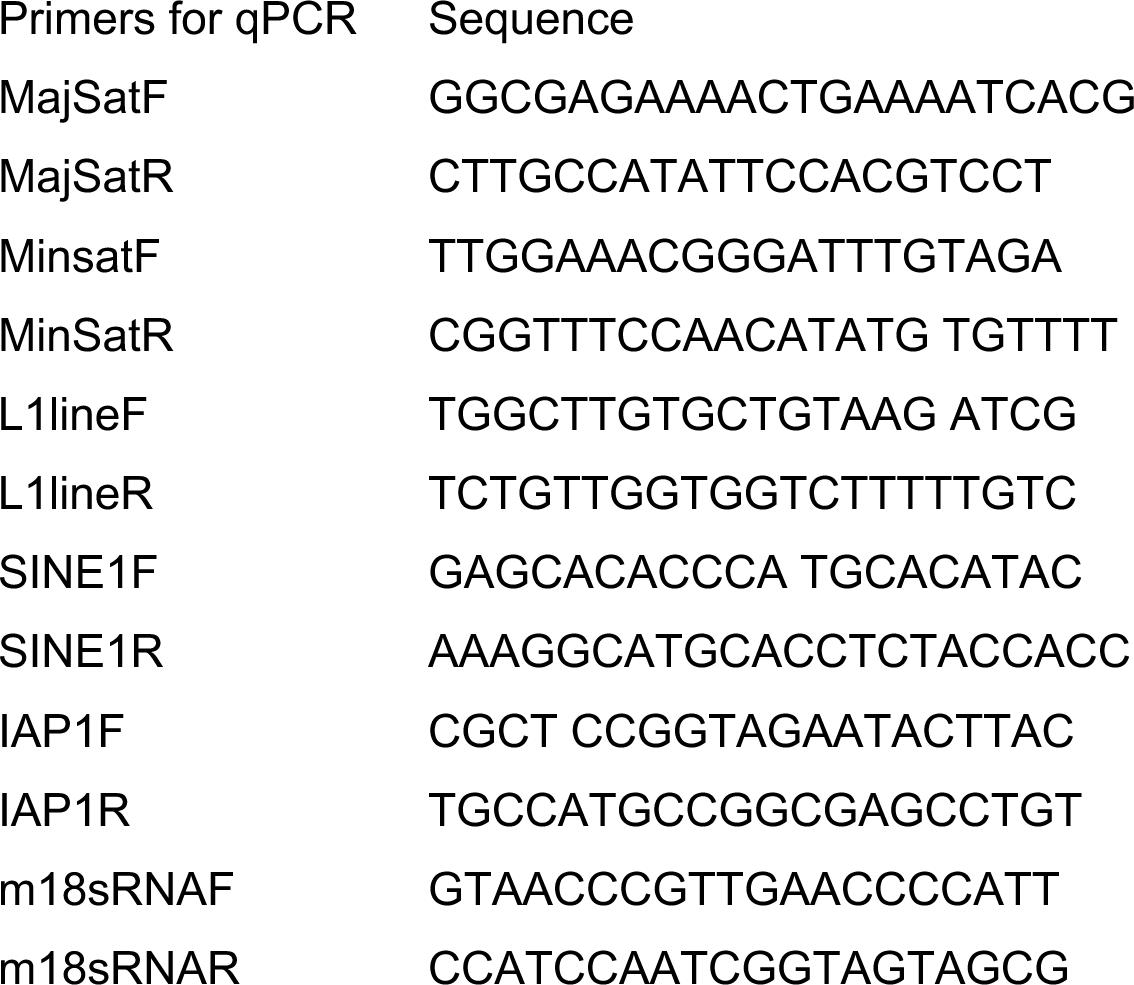

#### RNA-seq

Total RNA was isolated from CD11b^+^ cells and LK cells using RNeasy plus mini kit (Qiagen 74134). RNA libraries were prepared using NEB UltraII RNA library Prep kit for Illumina (NEB E7770L) according to the manufacturer’s instructions, and sequenced using Illumina NovaSeq 6000 to yield 30 million paired-end reads (50x50 bases) of per each RNA-seq library. For mice or human samples, reads were aligned against the mouse (mm10) or human (T2T) genomes respectively, using STAR (6), with the parameters -- outFilterMultimapNmax 1, --outSAMtype BAM SortedByCoordinate –sjdbOverhang 100 to analyze gene expression. HT-Seq (7) was used to quantify the gene expression levels using the options htseq-count -s yes, -r pos, -a 10. Normalization and differential expression analyses were performed using DESEQ2 (8), with the parameters fitType parametric, alpha 0.05 and using the Benjamini & Hochberg method. Differentially expressed genes (DEGs) between WT and mutant cells were defined as those with adjusted P value <0.05 and a fold change (log2FC) +/- 1. For visualization of the data, we generated tracks using Deeptools (9), with the option bamCoverage. All related plots were made using R-Studio and Integrative Genome Viewer (IGV) (10). To measure the expression of Transposable Elements (TEs), we used the TE-transcript package with the option -multi to quantify ambiguously mapped reads and then perform the differential expression (DE) analysis with DESEQ2 (FDR cutoff of p<0.05). Genes or TEs with less than 10 reads total were pre-filtered in all comparisons as an initial step. For TE analysis on mouse samples, we specifically used the mm10 repeatMasker version from TE-transcripts (11). The expression of interferon-stimulated genes (ISGs) were analyzed as previously described (12).

#### Cleavage under target and release using nuclease (CUT&RUN)

CUT&RUN was performed as previously described (13). All antibodies were used at 1:100 dilution. The spike-in CUT&RUN was performed with a 1:20 ratio of *Drosophila* S2 cells to mouse cells in each reaction (14). Briefly, cells were harvested as above and washed twice with PBS followed by two washes with wash buffer (20 mM HEPES pH 7.5, 150 mM NaCl, 0.5 mM spermidine, and 1× protease phosphatase inhibitor). Cells and spike-in *Drosophila* S2 cells were then incubated with activated Con A beads at room temperature for 20 min, followed by incubation with the primary antibody or IgG control overnight. Unbound antibody was removed by two washes with wash buffer supplemented with 0.025% digitonin, after which protein A/G–micrococcal nuclease (pA/G-MNase) was added at a final concentration of 700 ng/ml, and incubated for 1 hour at 4°C. Following incubation, the bound complexes were washed twice with Digitonin-wash buffer and placed in a cold metal block for 5 min to prechill the tubes. CaCl_2_ was added to activate MNase at a final concentration of 2 mM and incubated for 30 min in a cold metal block on ice. The digestion reaction was neutralized by the addition of an equal volume of 2× STOP buffer (200 mM NaCl, 20 mM EDTA, 4 mM EGTA, 50 μg/ml ribonuclease A, and 40 μg/ml glycogen) followed by incubation at 37°C for 10 min to release cleaved DNA fragments. The protein-DNA complex was collected by centrifugation at 16,000*g* for 5 min. Digested DNA was purified via spin column according to the manufacturer’s instructions (Zymo Research D5201). CUT&RUN DNA libraries for next-generation sequencing were generated using the NEBNext Ultra II DNA library Prep Kit following the manufacturer’s instructions (NEB 7645S). Libraries were pooled in equal quantities and sequenced using Illumina HiSeq 2500 (Illumina) to produce ∼30 million paired-end reads (50x50 bases) per library.

Paired-end reads were aligned against genome reference mm10 using Bowtie2 (v2.4.5) (15) using default parameters. Duplicated reads and unassembled contigs were removed using Picard. The number of reads aligning to mouse or Drosophila chromosomes were calculated using Samtools (v1.6) view -c -F 260. The ratio between the total number of reads mapping against mouse or Drosophila were used to estimate the normalization factors. bamCoverage was used to generate the bedGraph maps, applying the normalization factors. bedGraphToBigWig (UCSC) was used to generate the bigwig tracks. To remove the technical noise, IgG signals were removed from the CUT&RUN signals using bigwigCompare --operation subtract (9). Average tracks per condition were estimated by using BigwigMerge and considering the number of samples. To measure genome-wide changes, we used a 10kb approach, using bedtools map. CUT&RUN 10Kb windows data was intersected with Hi-C data to determine the presence of each region in A or B-compartments. All related plots were made using R-Studio and Integrative Genome Viewer (IGV).

#### Data Analysis and Statistical Methods

For quantitative real-time PCR, the mRNA level was determined using the Ct method with normalization to the housekeeping gene 18S ribosomal RNA. For all statistical tests, the exact “n” values can be found in figure legends. All quantitative data are presented as mean ± standard deviation from at least three samples or experiments unless otherwise stated. The significance was determined by two-tailed unpaired Student’s t-test, one-way ANOVA with Tukey’s multiple comparisons, and two-way ANOVA with Tukey’s multiple comparisons. Otherwise, methods are stated in figure legends. Statistical significance is shown as: *\*, P*< 0.5; *\*\*, P*< 0.01; and *\*\*\*, P*< 0.001. Statistical analyses were performed using RStudio and GraphPad PRISM software.

**Fig. S1.**
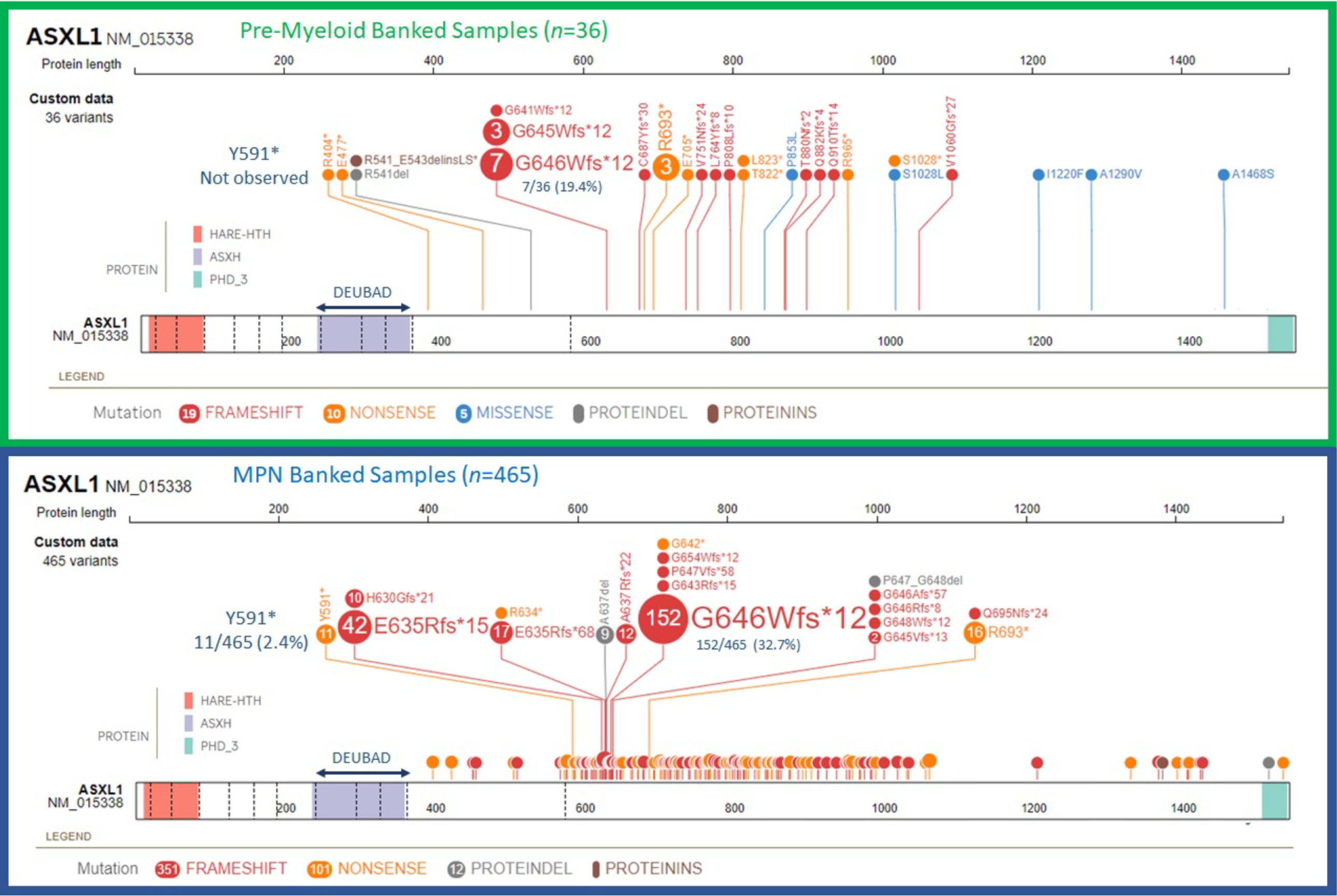
Spectrum of ASXL1 mutations observed in CH and CMML. The frameshift mutation c.1934dupG; p.G646WfsX12 (here termed G646fs) is by far the most common, both in pre-neoplastic CH (pre-myeloid samples, *top*) and in myeloproliferative neoplasms (MPN, *bottom*). This mutation yields a mutant ASXL1G646fs protein containing the first 646 amino acids of ASXL1 plus 12 additional amino acids not found in wildtype ASXL1. Also depicted is the Y591X mutation, which is represented only in MPN but encodes a biochemically better-behaved protein, all of whose amino acids are present in full-length ASXL1. All CH/CMML *ASXL1* mutations preserve the DEUBAD deubiquitinase adapter domain (aa 240-390). Most mutations are encoded in the large penultimate exon (exon 12) of *ASXL1*.

**Fig. S2.**
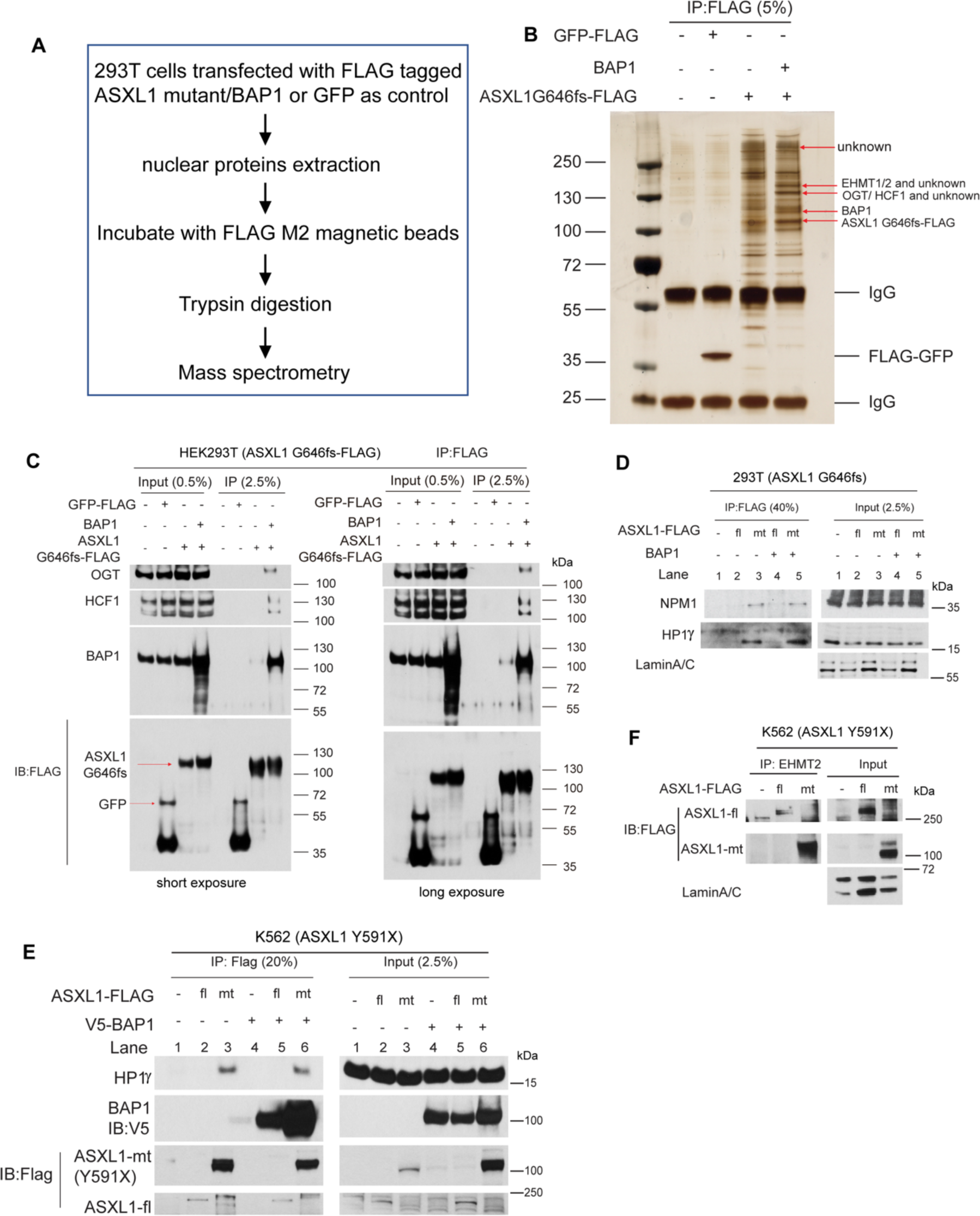
Mass spectrometry identifies novel heterochromatin-related proteins interacting with mutant ASXL1 G646fs. **A.** Workflow of immunoprecipitation followed by mass spectrometry (IP-MS) in HEK293T cells transfected with ASXL1G646fs-3xFLAG and BAP1. Cells expressing GFP were used as control. The migration positions of co-IP’d proteins are indicated. **B.** Silver staining of nuclear proteins co-immunoprecipitated with ASXL1G646fs-3xFLAG using anti-FLAG M2 magnetic beads. ASXL1G646fs-3xFLAG, BAP1, HCF1 and OGT are present in the immunoprecipitates. FLAG-tagged GFP is the control for IP. « - », cells transduced with empty vector. **C.** Western blot for OGT, HCF1 and BAP1 in immunoprecipitates of full-length ASXL1-3xFLAG or ASXL1G646Wfs-3xFLAG from HEK293T nuclear extracts. FLAG-tagged GFP is the control for IP. **D.** Verification of the interaction between mutant ASXL1G646Wfs and NPM1 or HP1γ in HEK293T cells. **E.** Immunoprecipitation in K562 cells constitutively expressing full-length ASXL1-3xFLAG and mutant ASXL1Y591X-3xFLAG shows that HP1γ preferentially interacts with mutant ASXL1. The K562 cell line carries the ASXL1 Y591X mutation on one allele. **F.** Endogenous EHMT2 pulls down full-length-3xFLAG and mutant ASXL1Y591X-3xFLAG from K562 cells stably expressing these proteins.

**Fig. S3.**
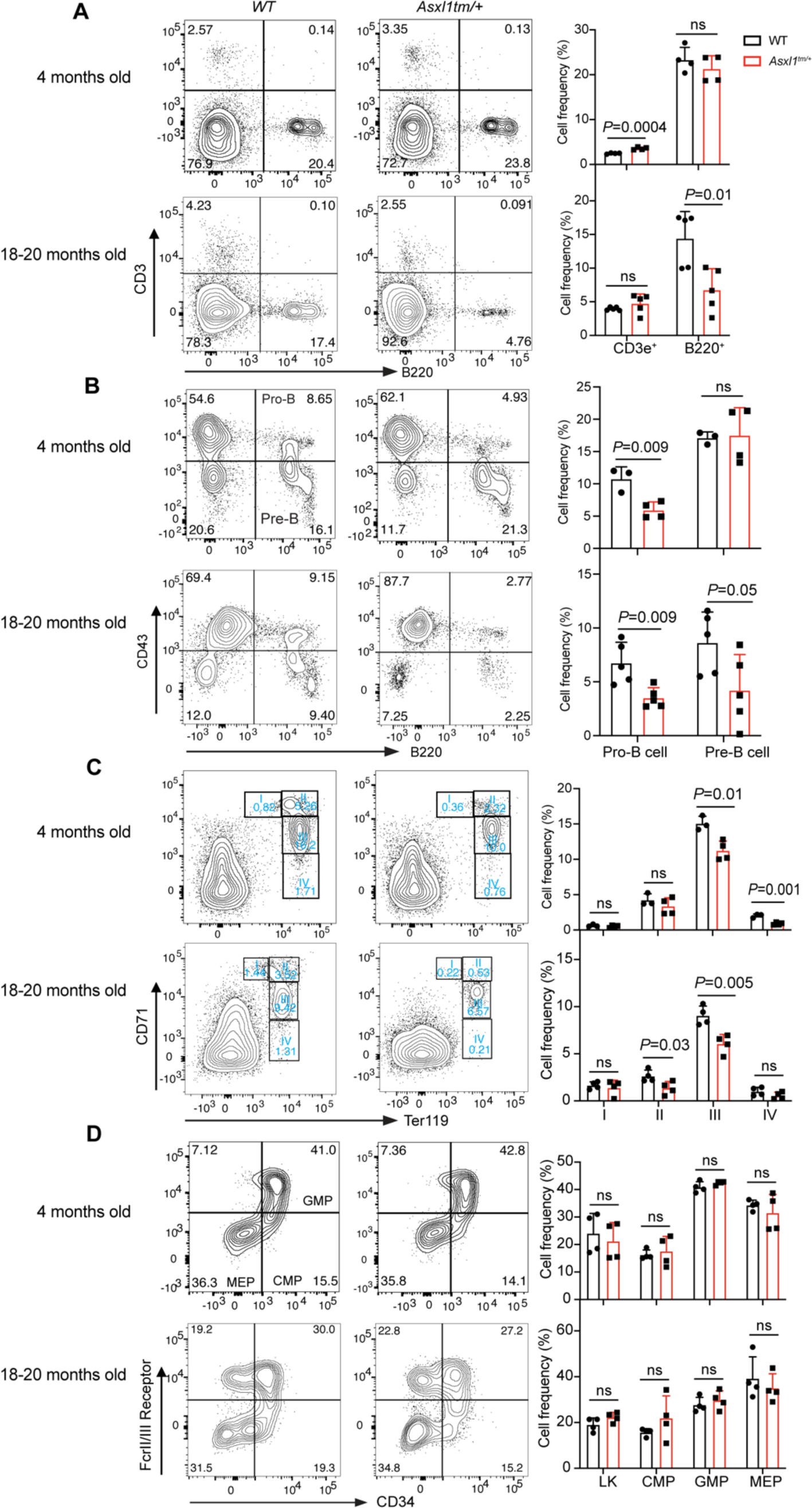
Hematopoietic development in young and old mice. **A.** Flow cytometry to assess T cell (CD3^+^) and B cell (B220^+^) lymphoid lineages in bone marrow of young (4 months, n=4 for each genotype) and old (18-20 months, n=5 for each genotype) WT and *Asxl1tm/+* mice. **B.** Flow cytometry and frequency of B cell lymphoid (B220/CD43) populations in the bone marrow of young (4 months, n=3 for WT, n=4 for *Asxl1tm/+*) and old (18-20 months, n=5 for each genotype) WT and *Asxl1tm/+* mice. **C.** Flow cytometry and frequency of erythroid (CD71/Ter-119) populations in bone marrow of young (4 months, n=3 for WT, n=4 for *Asxl1tm/+*) and old (18-20 months, n=4 for each genotype) WT and *Asxl1tm/+* mice. **D.** Flow cytometry of hematopoietic stem/progenitor cells (HSPC) in young (4 months, n=4 for each genotype) and old (18-20 months, n=4 for each genotype) WT and *Asxl1tm/+* mice. Cell populations were gated on Lineage^-^c-Kit^+^ Sca1^-^ (LK) cells. CMP, common multipotent progenitors; GMP, granulocyte-monocyte progenitors; MEP, megakaryocyte-erythrocyte progenitors. The summary graphs of flow cytometry are presented as mean ± SD. Student’s t-test is used to determine the *P* value.

**Fig. S4.**
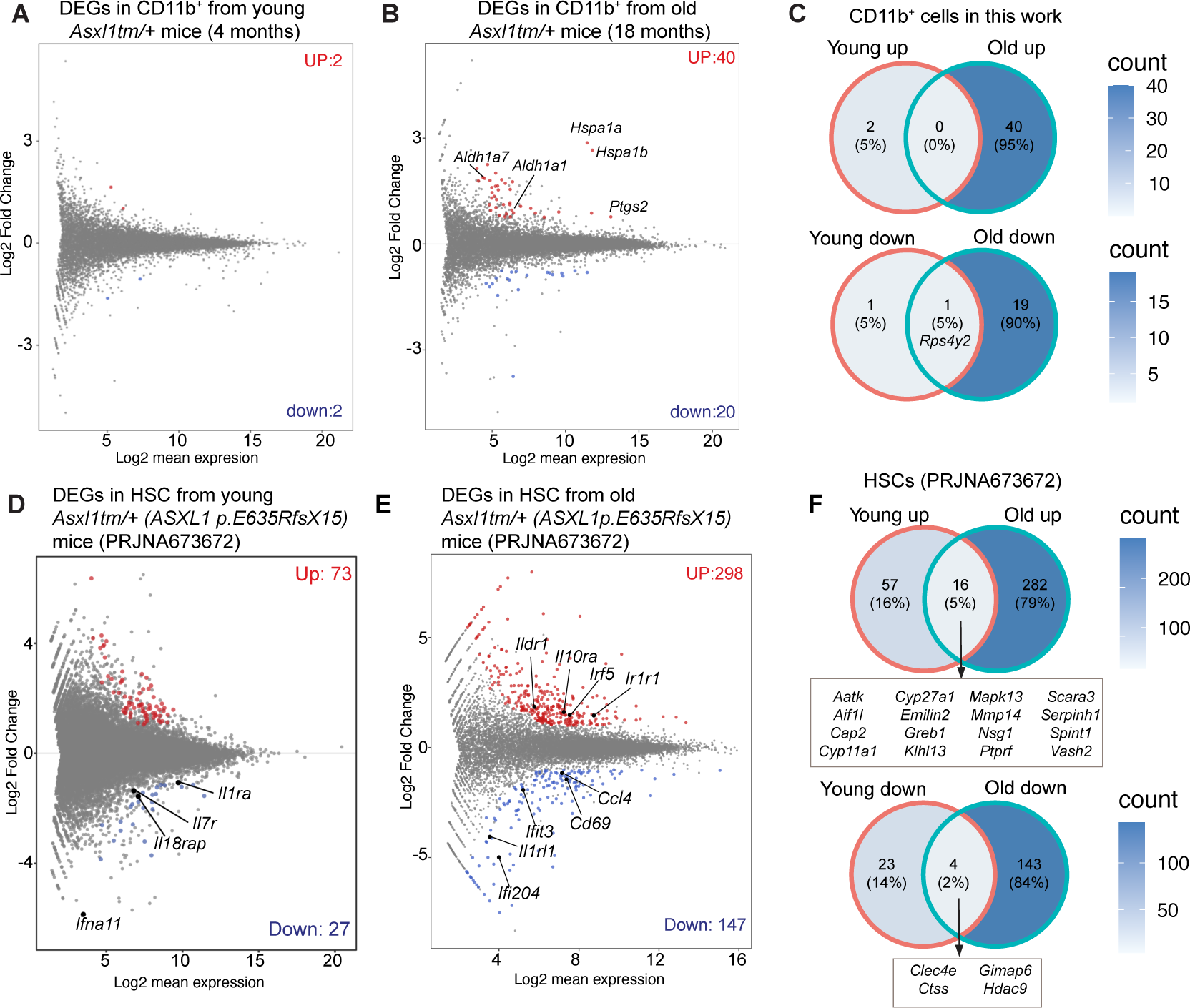
RNA-seq analysis of differentially expressed genes in CD11b^+^ cells and HSPCs from young and old *Asxl1tm/+* mice. **A.** MA plot of RNA-seq of CD11b^+^ cells from young *Asxl1tm/+* mice (4 months old). There were only 2 upregulated genes and 2 downregulated genes. **B.** Differentially expressed genes in CD11b^+^ cells from old *Asxl1tm/+* mice (18 months). Red dots represent genes expressed at high levels in *Asxl1tm/+* cells while blue dots represent downregulated genes. Some ISGs are upregulated, including *Aldh1a1*, *Aldh1a7*, *Hspa1a/b* and *Ptgs2*. **C.** Venn diagram of differentially expressed genes in CD11b^+^ cells from young and old *Asxl1tm/+* mice. No genes were upregulated in both young and old *Asxl1tm/+* CD11b^+^ cells, while only one gene, the *Rps4y2* gene encoding the ribosomal protein S4 Y-linked 2, is expressed at a low level in both young and old *Asxl1tm/+* CD11b^+^ cells. **D.** RNA-seq of genes in long-term hematopoietic stem cells (LT-HSCs) in young mice bearing the ASXL1 p.E635RfsX15 mutation. Reanalysis of RNA-seq data *PRJNA673672*. **E.** There were more differentially expressed genes in LT-HSC of old mice bearing the ASXL1 p.E635RfsX15 mutation. Reanalysis of RNA-seq data *PRJNA673672*. **F.** Overlap of differentially expressed genes in HSCs from young and old Asxl1 mutant mice (reanalysis of RNA-seq data, *PRJNA673672*). Only 16 genes showed high expression in both young and old *Asxl1tm/+* HSCs, while 4 genes were downregulated.

**Figure S5.**
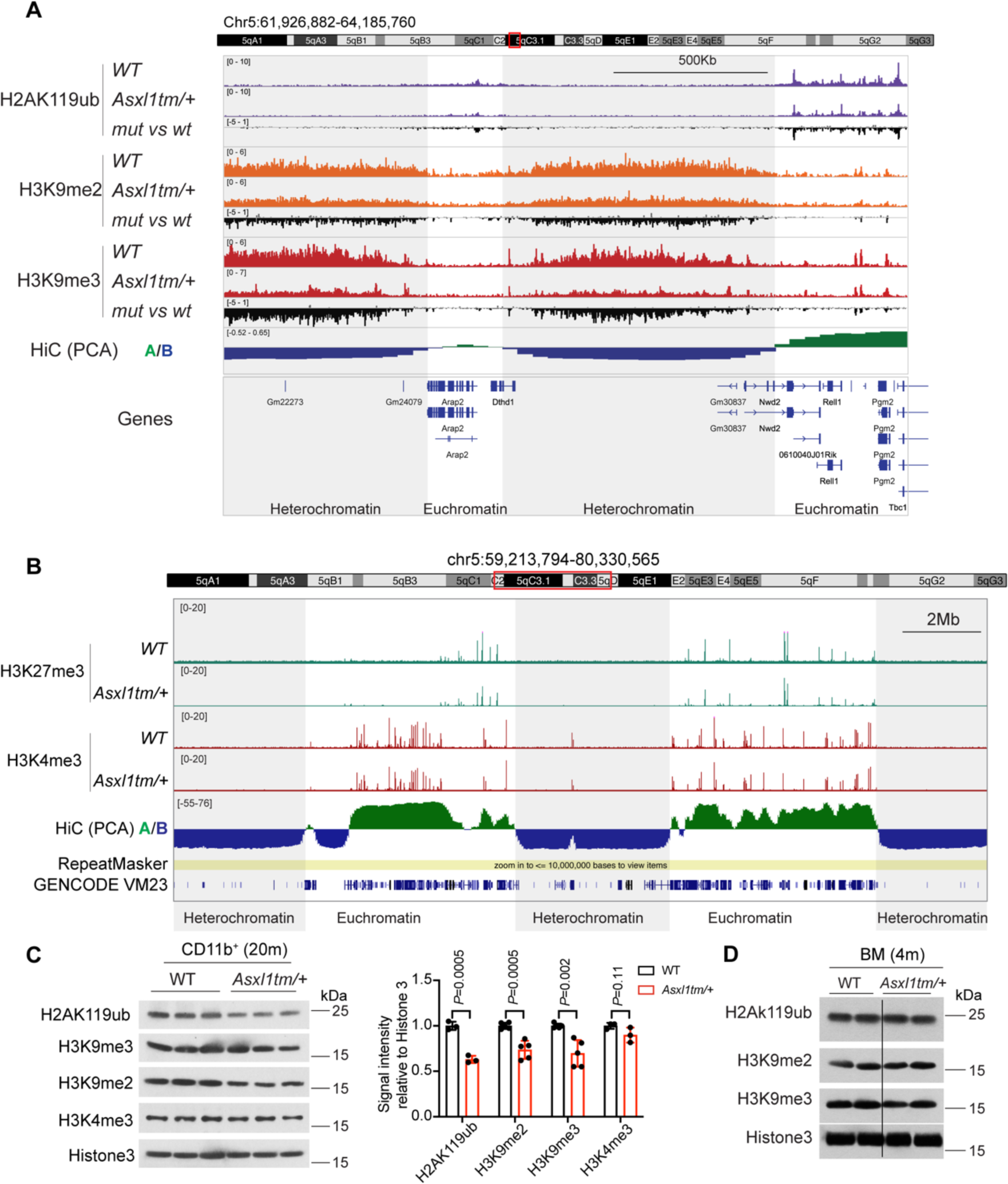
Altered histone modifications in *ASXL1tm/+* cells. **A.** A genome browser view of CUT&RUN data for H2AK119ub, H3K9me2 and H3K9me3 enrichment at the *Arap2* and neighboring genes in CD11b^+^ cells from 18-month-old *Asxl1tm/+* mice. *Drosophila* S2 cells were used as spike-ins for normalization. A ∼2.3 Mb region of chr. 5 is shown. Hi-C PC1 values were used to divide the genome into euchromatic (Hi-C A, positive PC1 values) and heterochromatic (Hi-C B) compartments. Compared to WT cells, *Asxl1tm/+* cells show global reductions of H2AK119Ub (*tracks 1-3)* in euchromatic regions (*track 10, green*) as well as H3K9me2/3 (*tracks 4-9*) in heterochromatin (*track 10, dark blue*); this is best appreciated from the difference tracks (*3, 6 and 9*). **B.** Genome browser view of H3K27me3 and H3K4me3 enrichment at euchromatin (*green*) and heterochromatin (*dark blue*) in CD11b^+^ cells from 18-month-old mice. H3K27me3 and H3K4me3 are present as discrete peaks in euchromatin. **C.** Western blots of acid-extracted histones for H2AK119ub, H3K9me2 and H3K9me3 in CD11b^+^ cells from old WT and *Asxl1tm/+* mice (20 months old) (*left panel*), and quantification of the relative signal intensity of histone modifications (normalized to histone 3) in CD11b^+^ cells (*right panel*). Student’s t-test suggests a significant difference in the levels of H2AK119Ub, H3K9me3 and H3K9me2 in bone marrow cells and CD11b^+^ cells of old *Asxl1tm/+* compared to age-matched WT mice, as well as a perceptible but not significant decrease in H3K4me3. **D.** Western blots of acid-extracted histones from bone marrow cells of young WT and *Asxl1tm/+* mice (4 months old) for H2K119ub, H3K9me2, and H3K9me3 modifications.

**Fig. S6.**
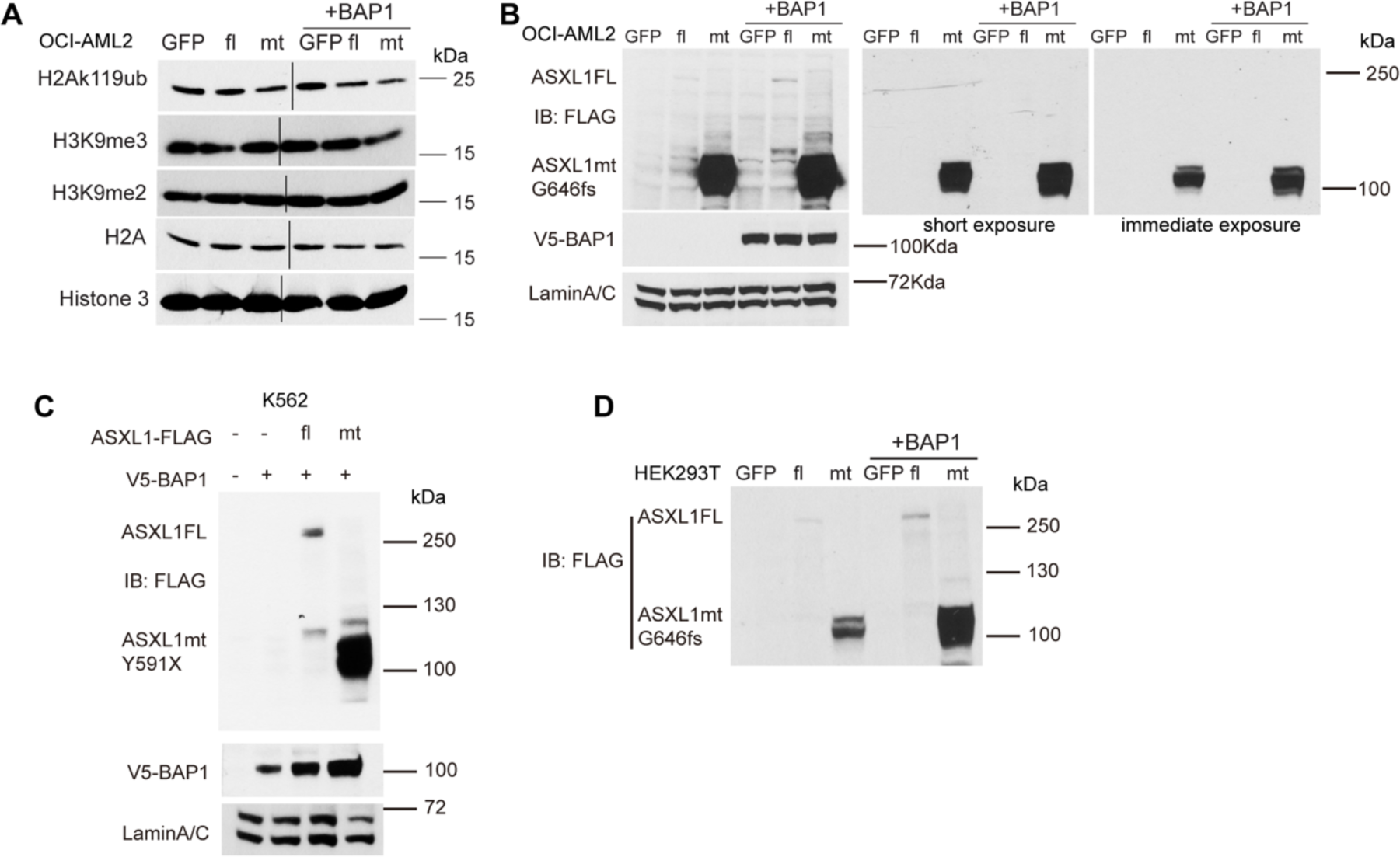
Expression level of ASXL1 full-length and mutant proteins in different cell types with ectopic expression. **A.** Western blots of acid-extracted histones from OCI-AML2 cells stably expressing full-length (fl) ASXL1-3xFLAG and mutant (mt) ASXL1G646fs-3xFLAG with or without BAP1 co-expression. Levels of the H2K119ub, H3K9me2, and H3K9me3 modifications were assessed, and showed a perceptible decrease in H2K119ub and H3K9me3 levels in cells co-expressing mutant ASXL1 and BAP1. **B.** Western blots of nuclear extracts from OCI-AML2 cells stably expressing full-length ASXL1-3xFLAG or ASXL1G646fs-3xFLAG, then lentivirally transduced with V5-tagged BAP1. Mutant ASXL1 is expressed at much higher levels compared to full-length ASXL1, and the levels of both proteins increase when BAP1 is co-expressed. The panels show the same gel exposed at different times. **C.** Western blot of nuclear extracts from K562 cells stably expressing V5-tagged BAP1 and transduced with either full-length (fl) ASXL1-3xFLAG or mutant (mt) ASXL1Y591X-3xFLAG. Mutant ASXL1Y591X is expressed at much higher levels than full-length ASXL1 in the presence of BAP1. **D.** Western blot of nuclear extracts from HEK293T cells transfected with full-length (fl) ASXL1-3xFLAG or ASXL1G646fs-3xFLAG, with or without V5 tagged BAP1. Mutant ASXL1 is expressed at much higher levels compared to full-length ASXL1, and the levels of both proteins increase when BAP1 is co-expressed.

**Fig. S7.**
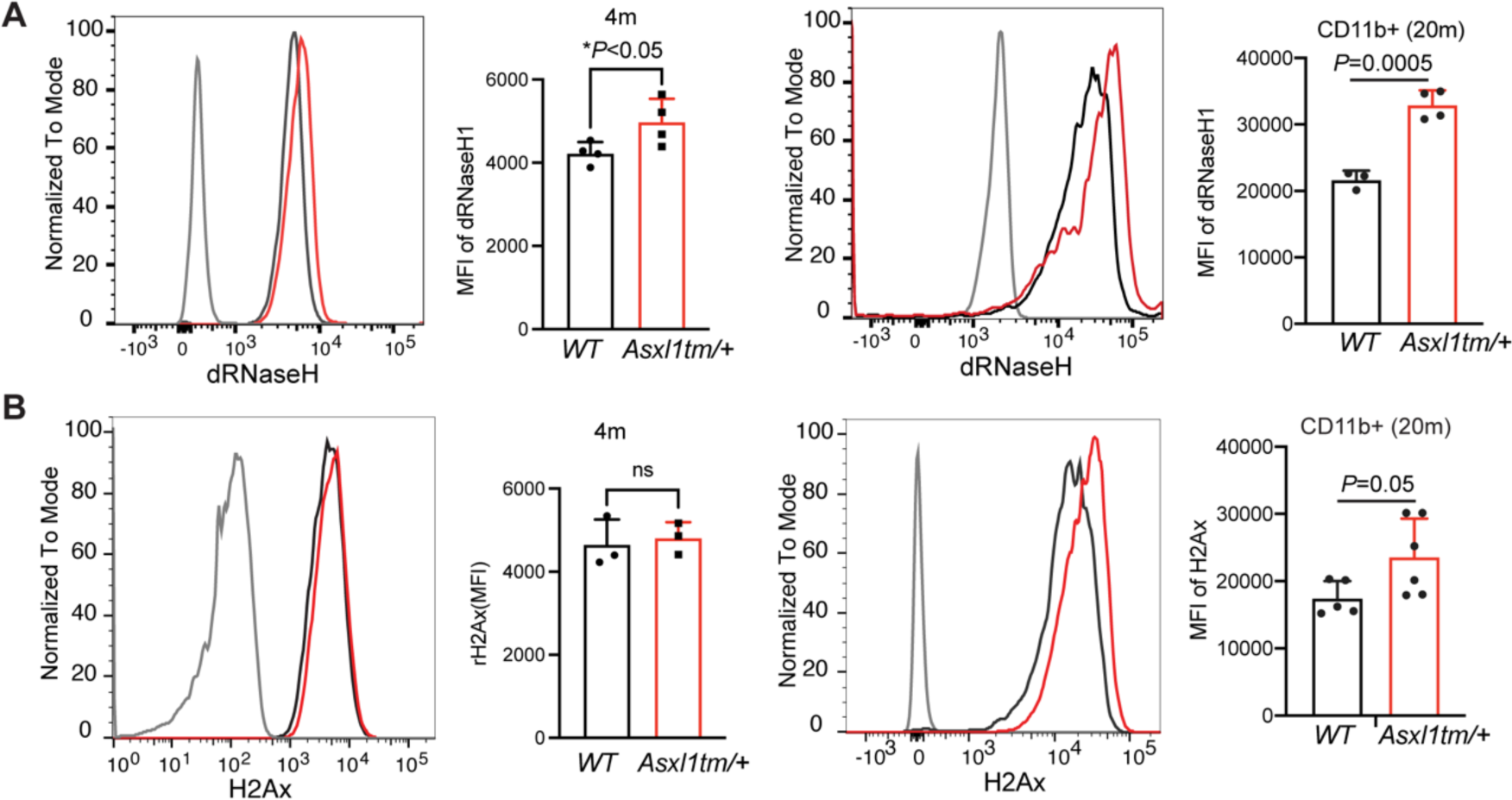
Increased R-loops and **γ**H2Ax in CD11b^+^ cells from old *Asxl1tm/+* mice. **A.** Frequency of R-loops, assessed by flow cytometry of permeabilized cells using V5-tagged catalytically-dead RNase H1, in CD11b^+^ cells from 4 months-old and 20 months-old WT and *Asxl1tm/+* mice (n=4). **B.** Flow cytometry and frequency of γH2Ax in CD11b^+^ cells from 4 months-old (n=4) and 20 months-old WT and *Asxl1tm/+* mice (n=5). The summary graphs of flow cytometry are presented as mean ± SD. Student’s t-test is used to determine the *P* value.

**Fig. S8.**
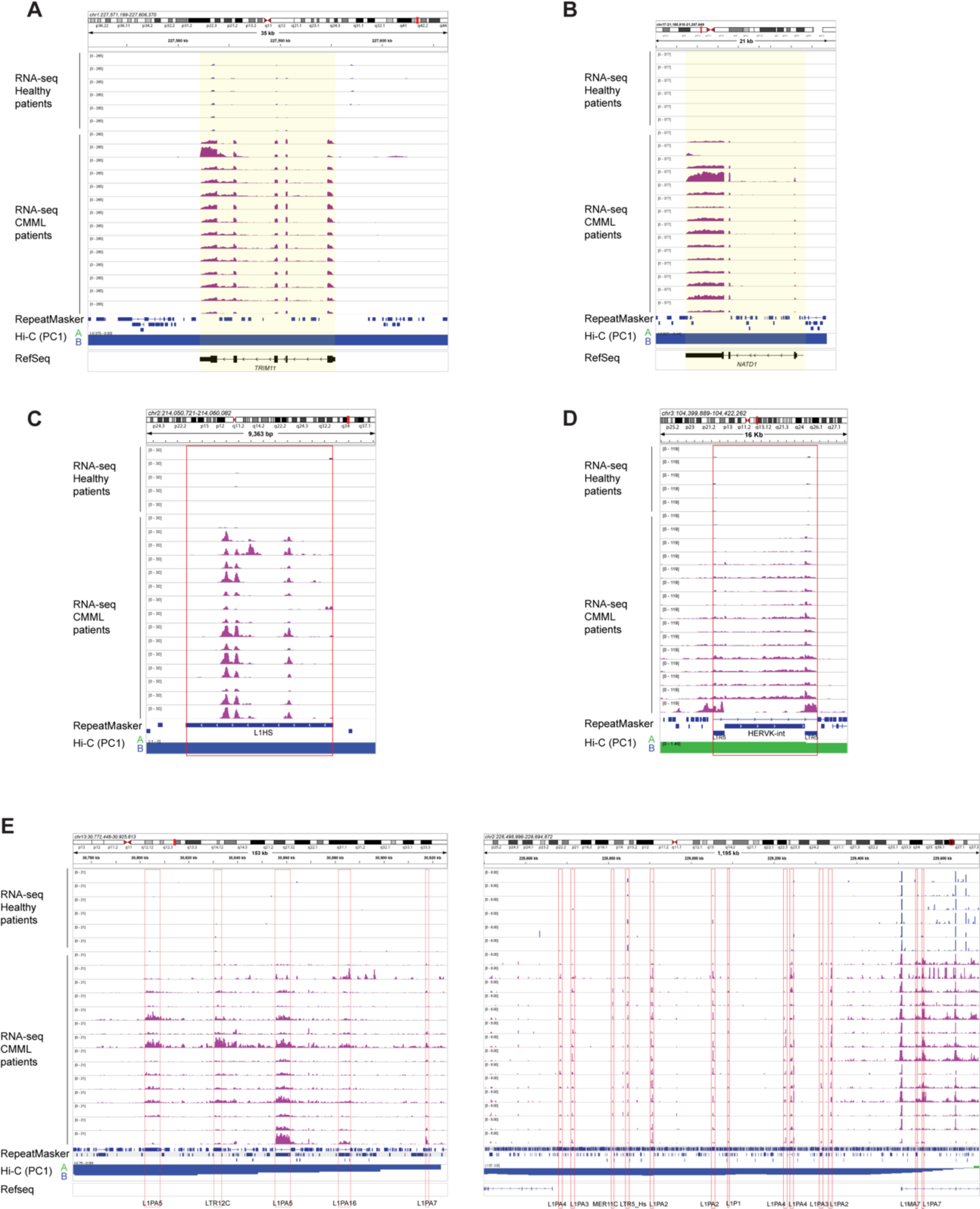
Increased expression of genes and TEs normally contained in heterochromatin in ASXL1-mutant CMML patients compared to healthy controls. **A-B.** Genome browser views showing upregulated genes at heterochromatin in *Asxl1*-mutant CMML (*bottom*, purple) samples compared to healthy controls (*top*, blue). **C-D**. Genome browser of unique L1HS (**C**) at heterochromatin and HERK-int (**D**) elements at euchromatin that are highly expressed in all *Asxl1*-mutant CMML samples. **E.** Clusters of TEs in heterochromatin that displayed higher expression in *ASXL1*-mutant CMML samples compared to healthy controls. Hi-C data from normal monocytes was used for chromatin compartmentalization. PC1 positive values (*green*) indicate euchromatin while negative values (*blue*) indicate heterochromatin.

**Fig. S9.**
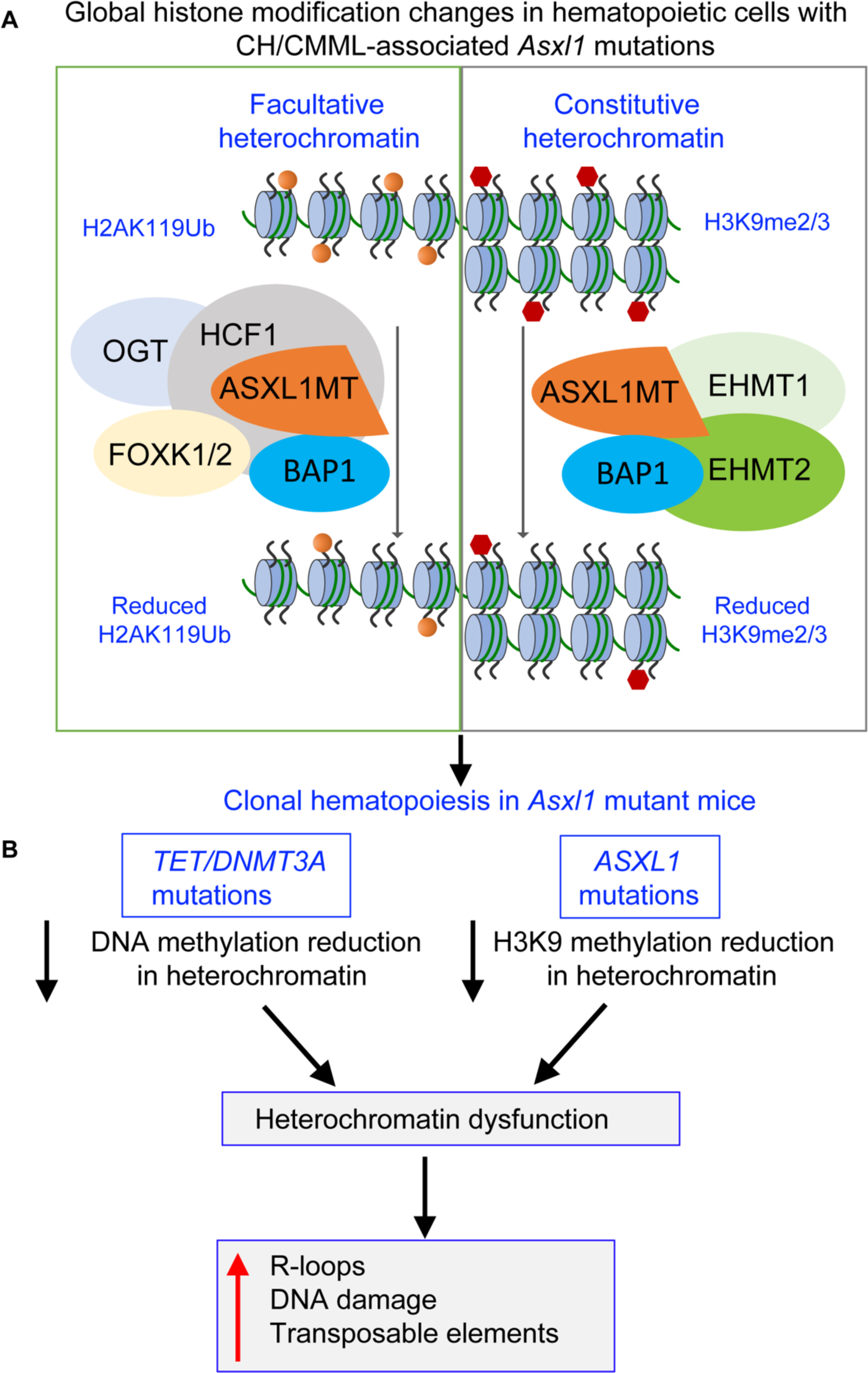
Working model for the role of mutant ASXL1. **A.** Two distinct ASXL1 protein complexes act in facultative and constitutive heterochromatin to cause global reduction of H2AK119Ub and H3K9me3 respectively. In *Asxl1*-mutant cells, C-terminally truncated mutant ASXL1 forms a canonical complex with BAP1, OGT, HCF1 and possibly FOXK1/2, which reduces levels of the H2AK119Ub modification present in both euchromatin and facultative heterochromatin. Our study reveals the existence of a new predicted complex, in which the mutant ASXL1-BAP1 heterodimer interacts with the EHMT1/2 H3K9me1/me2 methyltransferases, resulting indirectly in reduced levels of H3K9me2/me3 in constitutive heterochromatin. Decreases in both H2AK119Ub and H3K9me2/me3 are observed in clonal hematopoiesis caused by *Asxl1* mutations in our *Asxl1tm/+* mouse model. **B.** The three most frequent mutations in clonal hematopoiesis (in *TET2*, *DNMT3A* and *ASXL1*) compromise heterochromatin integrity. Loss-of-function mutations of *TET2* and *DNMT3A* result in decreased DNA methylation in heterochromatin. We show here that C-terminally truncated ASXL1 proteins also decrease H3K9me2/me3 levels in heterochromatin, through an indirect mechanism that remains to be defined. Thus, the top three mutations in clonal hematopoiesis all contribute to heterochromatin dysfunction.

